# Clinical and molecular characterisation of primary refractoriness to atezolizumab plus bevacizumab in patients with unresectable hepatocellular carcinoma

**DOI:** 10.1101/2025.07.08.663653

**Authors:** Pasquale Lombardi, Erik Ramon-Gil, Leonardo Brunetti, Giulia Francesca Manfredi, George Merces, Claudia Angela Maria Fulgenzi, Antonio D’Alessio, Aria Torkpour, Ciro Celsa, Bernardo Stefanini, Hannah Yang, Fionnuala Crowley, Thomas U. Marron, Anwaar Saeed, Matthias Pinter, Bernhard Scheiner, Yi-Hsiang Huang, Pei-Chang Lee, Naoshi Nishida, Ryan Po-Ting Lin, Andrea Dalbeni, Caterina Vivaldi, Gianluca Masi, Natascha Rohlen, Johann von Felden, Ahmed Kaseb, Peter R. Galle, Masatoshi Kudo, Wei-Fan Hsu, Lorenza Rimassa, Alessandro Parisi, Robin Kate Kelley, Hidenori Toyoda, Mario Pirisi, Rahim Falah Jabar, Mehrdad Rakaee, Giuseppe Cabibbo, Calogero Cammà, Fabio Piscaglia, Sohyun Hwang, Dong Jun Shin, Michael Li, Gennaro Daniele, Derek A. Mann, Hong Jae Chon, Jack Leslie, David J. Pinato

## Abstract

**Background:** Despite improved outcomes with atezolizumab plus bevacizumab (A+B) in hepatocellular carcinoma (HCC), primary refractoriness (PRef), characterised by early progression or short-lived disease stabilisation following treatment, remains a significant and poorly understood clinical challenge.

**Methods:** We analysed 1296 patients with HCC and Child-Pugh A liver cirrhosis treated with frontline A+B (AB-real) and validated findings in 645 trial participants recruited to IMbrave150 and GO30140. PRef was defined by Society for the Immunotherapy of Cancer (SITC) criteria as progressive disease in the first 6 months after treatment initiation. Patients who achieved complete response, partial response or stable disease for ≥ 6 months were classified as responders. We performed a multi-parametric analysis of pre-treatment tumour tissue including machine learning-based quantification of tumour-infiltrating lymphocytes, imaging mass cytometry and RNA sequencing (RNAseq) to evaluate differences in the tumour microenvironment (TME) of PRef versus responding patients. We employed conditional inference tree analyses to provide a hierarchical organisation of determinants of PRef.

**Results:** Among 677 AB-real and 378 trial patients evaluable by SITC criteria, PRef identified inferior median OS in comparison with responding patients (AB-real: 7.3 vs. 31.5 months, HR 3.7, 95%CI 2.8–8.5, p<0.001; Trials: 10.8 vs. NR, HR 4.6, 95%CI 3.3–6.3, p<0.001). PRef patients exhibited higher baseline systemic inflammation (neutrophil-to-lymphocyte ratio, NLR ≥3), a distinctively immunosuppressive TME enriched in CD163^+^ tumour-associated macrophages and a higher T^reg^/T^eff^ ratio. RNAseq of tumour tissue demonstrated lower intrinsic immunogenicity in PRef samples, characterised by repressed IFN-γ and T^eff^ signatures, with elevated myeloid infiltration. Conditional inference tree analysis identified IFN-γ signature downregulation combined with NLR ≥3 as the strongest contributor of PRef.

**Conclusions:** PRef to A+B identifies a distinct biological entity characterised by unopposed systemic inflammation, myeloid cell infiltration and T-cell depletion. Targeting myeloid-mediated immunosuppression, particularly in patients with low IFN-γ signature expression and elevated NLR might enhance responsiveness to A+B.

**Highlights:**

- Primary refractoriness to atezolizumab plus bevacizumab in hepatocellular carcinoma, as defined by SITC criteria, is associated with poor clinical outcomes.
- Tumour microenvironment profiling reveals an immunosuppressive phenotype characterized by high myeloid infiltration, reduced interferon-γ signalling, and T-cell depletion.
- The combination of systemic inflammation and low IFN-γ signature expression strongly predicts primary refractoriness and may inform therapeutic decision-making.

## Introduction

The prognostic outlook of patients with unresectable-advanced hepatocellular carcinoma (HCC) has remarkably improved in the last years due to the introduction of immune checkpoint inhibitors (ICI), which have shifted the expected median overall survival (OS) from approximately 10-13 months with sorafenib to up to 19.2 months with the combination of atezolizumab plus bevacizumab (A+B), the first immunotherapy combination to be introduced in clinical practice ^1–5^.

Whilst survival improvement from the use of Programmed Cell Death-1 (PD-1)/PD-1 ligand (PD-L1) inhibitor-containing combinations has been amply validated across several combination studies with Vascular Endothelial Growth Factor (VEGF), tyrosine kinase inhibitors (TKI) and Cytotoxic T-lymphocyte antigen-4 (CTLA-4) inhibitors ^6, 7^, the proportion of patients that derive a measurable response remains confined to a minority ranging from 20 to 36% ^6^.

Although achieving a response is not a universal pre-requisite to securing an overall survival benefit from all ICI combinations ^2, 4^, disease progression on anti-cancer immunotherapy remains invariably associated with a worse prognosis in HCC ^2, 4, 8, 9^. Therapeutic resistance to ICI remains an area of high unmet need, with patients experiencing either adaptive resistance following an initial response or primary resistance characterised by early disease progression on treatment. In solid tumours, primary refractoriness (PRef) to immunotherapy coincides with low intrinsic immunogenicity of the tumour, which can manifest itself through altered expression of tumour-associated antigens or lower neo-antigen load, defective epitope presentation and enrichment of immune co-inhibitory signals and immunosuppressive immune cells within the tumour microenvironment ^10^. These characteristics are uniquely different from those driving adaptive resistance to ICI and bear a comparatively worse contribution to the prognosis of immunotherapy recipients ^10^. Acknowledging the fundamental differences in the immunobiology underlying the various patterns of resistance, the Society for Immunotherapy of Cancer (SITC) has developed consensus recommendations for defining resistance to ICI based on observed responses to treatment, providing a standardised framework to reproducibly identify PRef tumours in the clinic ^11^.

Despite their demonstrated clinical utility in clinical research and routine practice, these criteria have not been validated in HCC, a disease where prognosis is not only dictated by tumour progression but uniquely influenced by concomitant liver dysfunction ^12, 13^. In addition, whilst efforts have been addressed to identify predictors of response to A+B, little attention has been dedicated to the phenotypic characterisation of those patients who will not derive upfront clinical benefit from A+B ^14, 15^.

In this study, we aimed to portray the clinical course of HCC patients who are primarily refractory to A+B. Using clinical data and pre-treatment HCC tissue samples from a large observational study and two landmark clinical trials in unresectable/advanced HCC, we sought to validate PRef to A+B as defined by SITC criteria and investigate the biological characteristics underlying PRef to identify prognostic factors and potential therapeutic targets to overcome primary resistance.

## Methods

### Study design and clinical outcomes

Patient disposition across studied cohorts is summarized in ***Figure 1***. The study design and methods for AB-real ^8^, GO30140 ^16^ and IMbrave150 have been described previously^1^.

**Figure 1.**
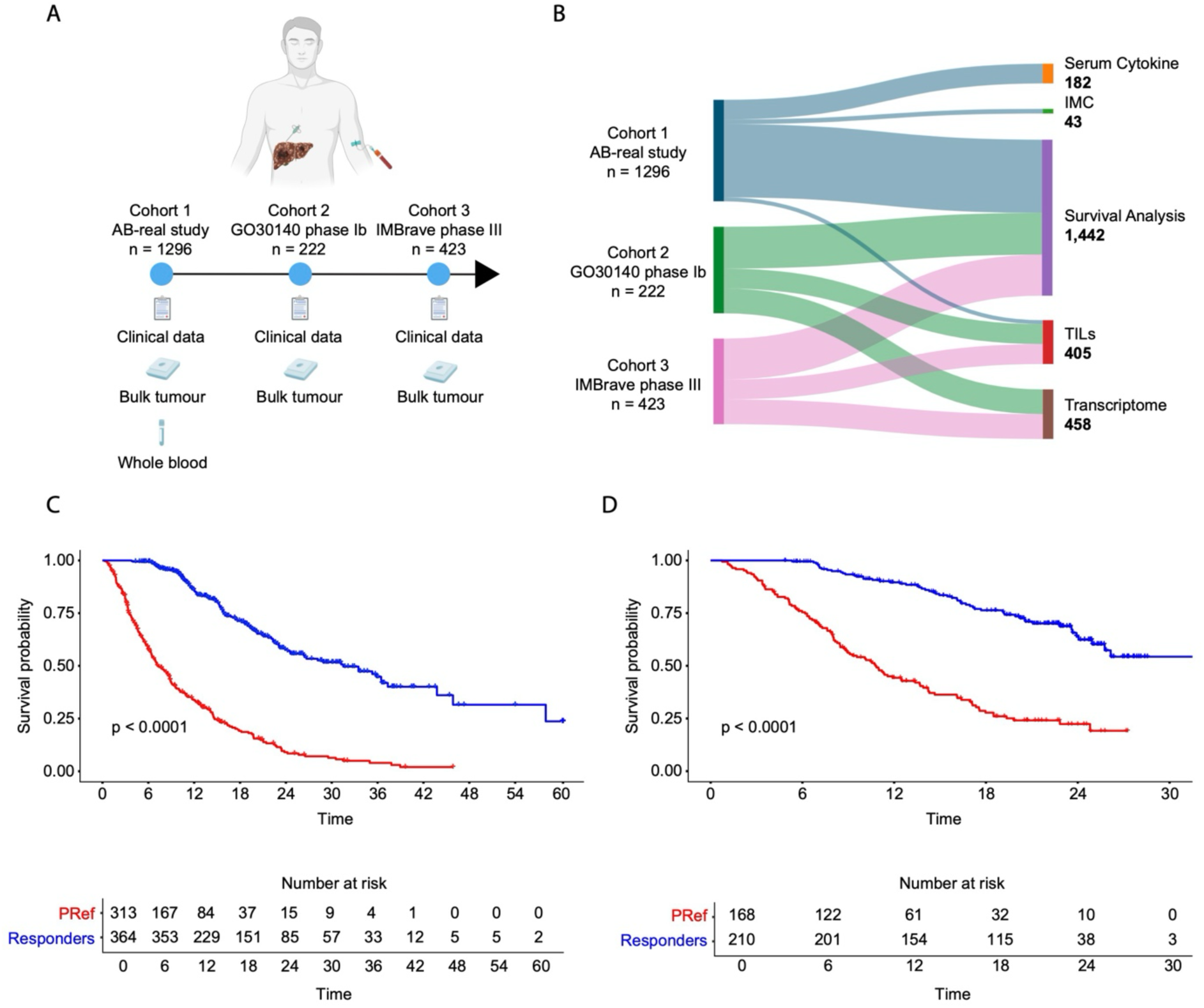
Study overview. (**A**) Overview of the three cohorts analysed, comprising 1296, 222 and 423 patients, respectively. (**B**) Sample utilisation from each cohort. (**C**-**D**) Kaplan-Meier curves of OS in Cohort 1 and Cohorts 2+3, stratified by PRef and Responders. AB, Atezolizumab + Bevacizumab; IMC, Imaging Mass Cytometry; TILs, Tumour-infiltrating lymphocytes; OS, Overall Survival; PRef, Primary Refractoriness.

The SITC consensus criteria were applied to identify primary resistance or refractoriness: PRef was defined as the achievement of progressive disease or stable disease lasting <6 months as best overall response to ICI in all cohorts ^11^. Responders were defined as patients who achieved a complete response (CR), a partial response (PR) or stable disease lasting ≥6 months as best overall response to ICI in all cohorts. Radiological response to treatment was assessed according to RECIST criteria v1.1. Treatment duration was calculated from the date of the first dose of A+B until discontinuation. OS was defined as the time from the first A+B dose of the treatment until death or last follow-up. Details on study design, patient assessments and response outcomes are described in ***Supplementary Methods*.**

### Machine-learning approach for tumour infiltrating lymphocytes (TILs) identification and quantification

We applied a supervised machine learning (ML) algorithm to 37 and 184 H&E of Cohort 1 and Cohort 2-3, respectively, developed using QuPath (v.0.5.0) and followed the previously described procedure with some modifications ^17, 18^ ***(Figure 1)***. A detailed description of model development, training and quality verification can be found in the ***Supplementary Methods*.**

### Imaging Mass Cytometry (IMC)

Antibodies against human antigens were previously validated by immunohistochemistry (IHC), conjugated to IMC-compatible metal isotopes using the MaxPar X8 kit (Standard Biotools - 201300). Forty-three clinically annotated pre-treatment FFPE biopsy specimens were incubated with three rounds of Xylene, followed by decreasing concentrations of ethanol. Antigen retrieval was performed using Tris-EDTA (Ph9) at 98°C and tissues were blocked upon incubation with 3% Bovine Serum Albumin (ThermoFisher - A7906-100G) in PBS. Then, slides were incubated with the metal-conjugated antibody cocktail (***Supplementary Table 1***) overnight at 4°C. The next day, the excess of antibody was washed with TBST 1X and PBS, and nuclei were stained using the Cell-ID™ Intercalator-Ir—125 µM (Standard Biotools - 201192A). Finally, tissues were left drying at room temperature, ready to be ablated using the Hyperion imaging device (Standard Biotools).

Once samples were imaged, a visual QC was performed to ensure good staining quality of the regions of interest (ROIs) used for the downstream analysis. A total of 441 ROIs were used for data generation. Data was exported in .TIFF format and cell segmentation was performed following the OPTIMAL pipeline ^19^. Single cells were clustered in cell types using the FlowSOM algorithm from FCS Express (Lic. no 75616), and then exported for neighbour joining analysis to generate the cell–cell interaction data. Cell-cell interaction analysis was performed as previously described^20^.

### Serum cytokines quantification

Peripheral blood samples were obtained immediately before the first administration of A+B in 182 patients of cohort 1 ***(Figure 1*)**. Following centrifugation at 1,000×g for 5 minutes; serum samples were stored at -80 °C. Serum cytokine concentrations were quantified using a cytometric bead array (560484, BD Biosciences) according to the manufacturer’s instructions.

In brief, the capture beads were incubated with serum samples and the detection reagent for 3 hours at room temperature. After several washes, cytokine-bead complexes were measured using a flow cytometer (Beckman Coulter), and the data were analysed using the FlowJo software (Tree Star Inc.).

### Tissue gene expression analysis

Differential gene expression analysis was performed with DESeq2, comparing responders and PRef. Significant differential expression was defined as an absolute log₂ fold-change of >1 and an adjusted *P* < 0.05.

xCell deconvolution analysis was applied to baseline RNA-seq data, and cell type enrichment scores were compared between PRef and responders using Student’s t-test with Benjamini– Hochberg correction.

We calculated gene signature scores for each sample as the arithmetic mean of log2(CPM) expression of all genes in a given signature (***Supplementary Table 2***). Gene set enrichment analysis (GSEA) was performed using the MultiGSEA package v.1.1.99 with default parameters, and the Hallmark gene set collection from the molecular signatures database collection (MSigDB) and the CAMERA method for multiple hypothesis testing ^21^.

## Statistical analysis

Statistical analyses were performed using R v.4.3.1. Survival curves were estimated using the Kaplan-Meier method and compared with the log-rank test. Hazard ratios (HR) with 95% CIs and p values were determined using a stratified Cox proportional hazards model and log-likelihood test. The proportional hazards assumption was validated using Schoenfeld residuals.

Multivariate adjustment included key clinical and laboratory variables, detailed in the ***Supplementary Methods***. To assess the association between SITC-defined response and survival outcomes while minimizing immortal time bias, we employed a time-dependent Cox proportional hazards model. Response status was treated as a time-varying variable, with patients initially classified as PRef. Those who remained progression-free beyond six months were reclassified as responders, while others remained PRef. This approach improved the accuracy of hazard estimates and prevented artificial survival advantages.

To explore determinants of PRef, we applied a conditional inference tree (CTree) analysis using the party R package, visualising results with Sankey diagrams. The biomarker-evaluable population was analysed separately in Cohorts 2 and 3 to mitigate batch effects in RNA-seq data but was combined for CTree analysis, where RNA signatures were reduced to categorical variables. Detailed methods in the ***Supplementary Methods***.

## Results

### Patient Cohorts

Of the 3 parallel patient cohorts derived from prospective observational and randomized clinical studies (***Figure 1A***), Cohort 1 consisted of 1296 patients included in the prospectively maintained database AB-real, a subgroup of whom had pre-treatment tissue available for IMC (n=43) and serum available for cytokine/chemokine quantification (n=182). Cohorts 2 and 3 included 645 patients treated in the GO 30140 (n=222) phase I and IMbrave 150 (n=423) phase III clinical trials, respectively, with bulk transcriptome data and TIL quantification (n=229 and n=184 respectively) (***Figure 1B***). After removing patients who did not meet inclusion criteria, 677 and 378 patients were evaluated for SITC criteria for cohorts 1 and 2-3, respectively (***Supplementary Figure 1***).

### Primary refractoriness to A+B is associated with poor oncological outcomes in unresectable HCC

We first assessed the relationship between PRef and OS and used Cohort 1 as a training set and Cohort 2-3 as validation.

Among the 677 patients in Cohort 1, 313 (46%) were classified as PRef in Cohort 1 due to PD as first response (220, 70.3%) or SD <6 months (93, 29.3%). In Cohorts 2-3, 168 (44%) patients were identified as PRef due to PD (91, 54.2%) or SD<6 months (77, 45.8%).

Across Cohorts, PRef patients had a higher percentage of Barcelona Clinic Liver Cancer (BCLC) stage C disease (Cohort 1: 80% vs. 72%, *p*=0.0017; Cohort 2-3: 92% vs. 79%, *p=*0.001), and higher NLR ≥3 (Cohort 1: 60% vs. 37%, *p*<0.001; Cohort 2-3: 58% vs. 34%, *p*<0.001) compared to responders. Clinical characteristics and outcomes are summarised in ***Supplementary Tables 3-4*** and ***Supplementary Results***.

After a median follow-up of 20.9 (95% CI 19.8–22.0) (Cohort 1) and 14.3 (95% CI 8.1–21.1) months (Cohort 2-3), 308 (45.9%) patients and 207 patients (54.8%) were alive at the data cut-off in Cohort 1 and Cohort 2-3, respectively. Median OS was significantly worse in PRef vs. responders (Cohort 1: 7.3 vs. 31.5 months, HR 3.7, 95%CI 2.8–8.5, p<0.0001; Cohort 2-3: 10.8 vs. NR, HR 4.6, 95%CI 3.3–6.3, p<0.0001, ***Figure 1C-D***). To address the risk of immortal time bias, we performed time-dependent multivariate models, which confirmed PRef status to be associated with the highest risk of mortality across Cohorts (Cohort 1: HR 4.0, 95% CI 2.8– 5.5 p<0.001; Cohorts 2-3: HR 4.9, 95% CI 3.6–6.7, p<0.001) alongside NLR ≥3, α-fetoprotein (AFP) and Albumin-Bilirubin (ALBI) grade (***Supplementary Table 5***).

To further investigate the clinical characteristics of PRef, the NLR was chosen as a marker of systemic inflammation^22, 23^. PRef patients exhibited a higher baseline inflammatory status compared to responders as measured by median NLR of 3.6 vs. 2.7 in Cohort 1 and 3.3 vs 2.4 in Cohort 2-3 (p<0.001). The proportion of patients with NLR ≥3 was also higher across PRef vs responders in Cohort 1 (63% vs 40%, p<0.001) and Cohorts 2-3 (65% vs. 35%, p<0.001, ***Supplementary Tables 3-4***).

### PRef is independent of pre-treatment TIL density

We sought to determine whether PRef patients harboured a differentially immunogenic tumour microenvironment compared to responders and first investigated differences in tumour-infiltrating lymphocytes (TILs) density across groups, electing TILs as *bona fide* markers of spontaneously immunogenic tumours ^24^. We applied a pre-optimised ML algorithm to evaluate the differential density of TILs in a total of 37 patients treated with A+B in Cohort 1 and 184 patients accrued in Cohort 2-3 who had digitally available H&E slides. Median cell density was 141 (IQR 419) and 417 (IQR 494) cell/mm^2^ of assayed tissue in Cohort 1 and Cohort 2-3 respectively and not different in PRef compared to responders (Cohort 1: 138 vs 169 cell/mm2, p=0.74: Cohort 2-3 442 vs 399 cell/mm^2^, p=0.57, ***Figure 2A-B***).

**Figure 2.**
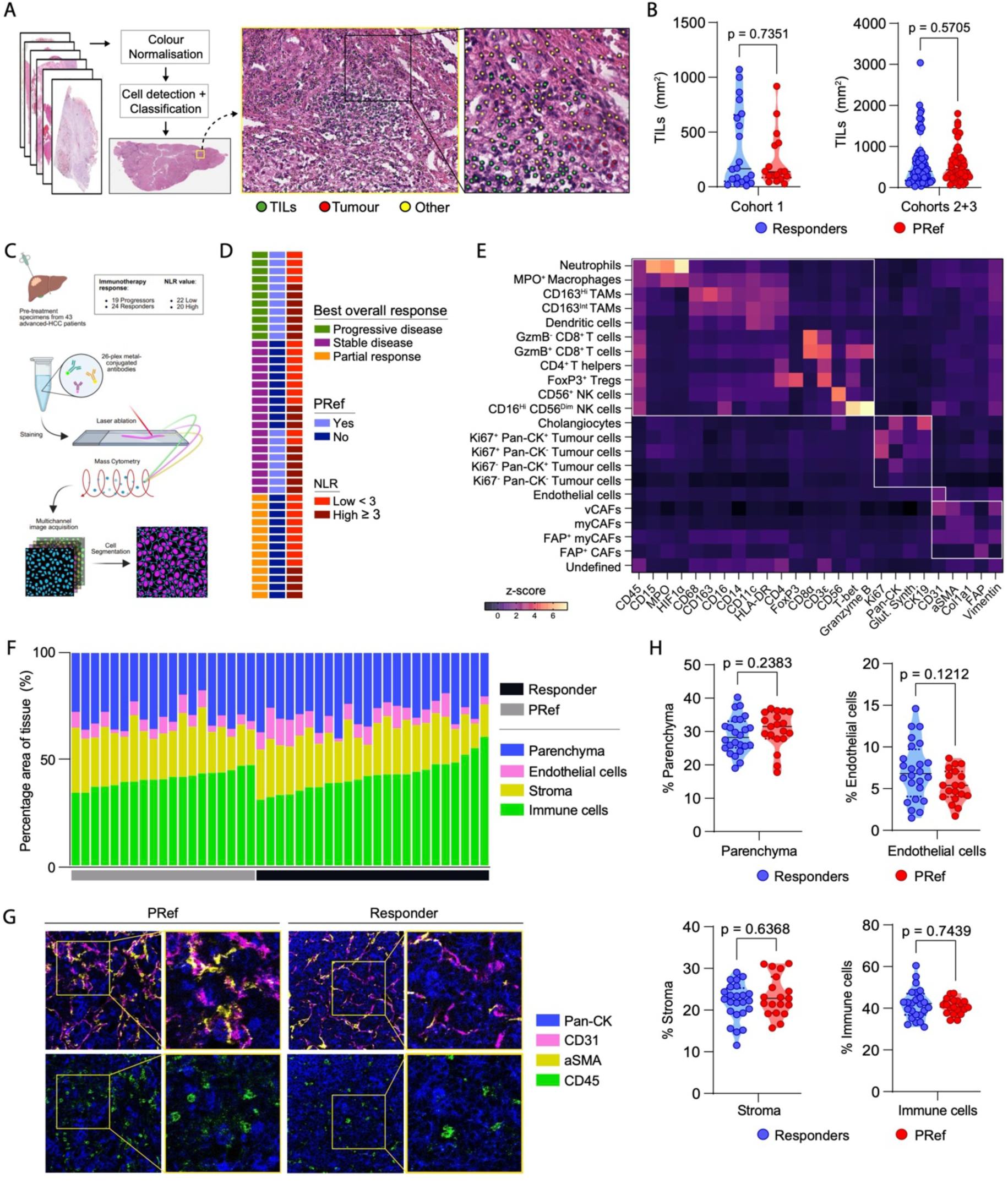
Decoding the HCC TME of PRef and Responders to Atezolizumab plus Bevacizumab. (**A**) Schematic of the machine learning-based TILs quantification. (**B**) TILs counts between PRef and Responders in Cohort 1 and 2. (**C**-**D**) IMC cartoon and clinical information of the cohort for response, PRef and NLR. (E) Identity HeatMap showing the cell types identified in the HCC TME by IMC. (F-H) IMC Data and representative figures for the comparison of whole Parenchyma, Endothelia, Stroma and Immunity between PRef and Responders. CD, Cluster of Differentiation; TME, Tumour Microenvironment; NLR, Neutrophil to Lymphocyte Ratio; MPO, Myeloperoxidase; TAMs, Tumour-associated macrophages; GzmB, Granzyme B; FoxP3, Forkhead box P3; NK, Natural Killer; PanCK, Pan-Cytokeratin; CAFs, Cancer-associated Fibroblasts; vCAFs, Vascular CAFs; myCAFs, Myofibroblast CAFs; αSMA, Alpha-smooth muscle actin; FAP, Fibroblast Activation Protein.

### Primary refractoriness to A+B is associated with a unique immune composition of the HCC TME

We next performed in-depth spatial immune phenotyping of the tumour microenvironment to characterise the mechanisms underpinning PRef to A+B. We performed highly multiplexed IMC in a subset of Cohort 1 patients with pre-treatment FFPE tissue available for analysis (n=43) (***Figure 2C***). In total, we acquired 441 high-dimensional histopathology images representing 43 patients with HCC; 19 PRef and 24 responders (***Figure 2D***). Images were segmented into 291,642 total cells, clustered and then annotated using the OPTIMAL IMC analysis pipeline ^19^. Using this approach we classified multiple tumour cell populations, various stromal and vascular cell types as well as 11 individual immune cell populations representing a range of lymphocyte and myeloid populations (***Figure 2E***). Quantification of these major cellular compartments showed no difference in the number of tumour, stromal, endothelial or immune cells in PRef compared to responders (***Figure 2F-H***). Therefore, a deeper and more detailed analysis of cell compartments was performed.

When we compared the abundance of individual cell types, we found that CD163^+^ tumour-associated macrophages (CD163^+^ TAMs), encompassing the entire spectrum of CD163 high (CD163^Hi^) and CD163 intermediate (CD163^Int^) expression, were enriched in PRef patients, being the CD163^Int^ TAMs the individual cell subpopulation more strongly associated with PRef (p<0.0001). These were followed by vascular cancer-associated fibroblasts (vCAFs) (p<0.05). Moreover, although not statistically significant, anti-tumour entities like CD8^+^ T-cells, CD4^+^ T helpers, dendritic cells and CD16^hi^ CD56^dim^ NK cells were enriched in responders, whereas pro-tumour cell types such as neutrophils, CD4^+^FoxP3^+^ T regulatory cells (FoxP3^+^ T^reg^) and a diversity of other CAF subtypes were enriched in PRef (***Figure 3A-B***, ***Supplementary Figure 2B***).

**Figure 3.**
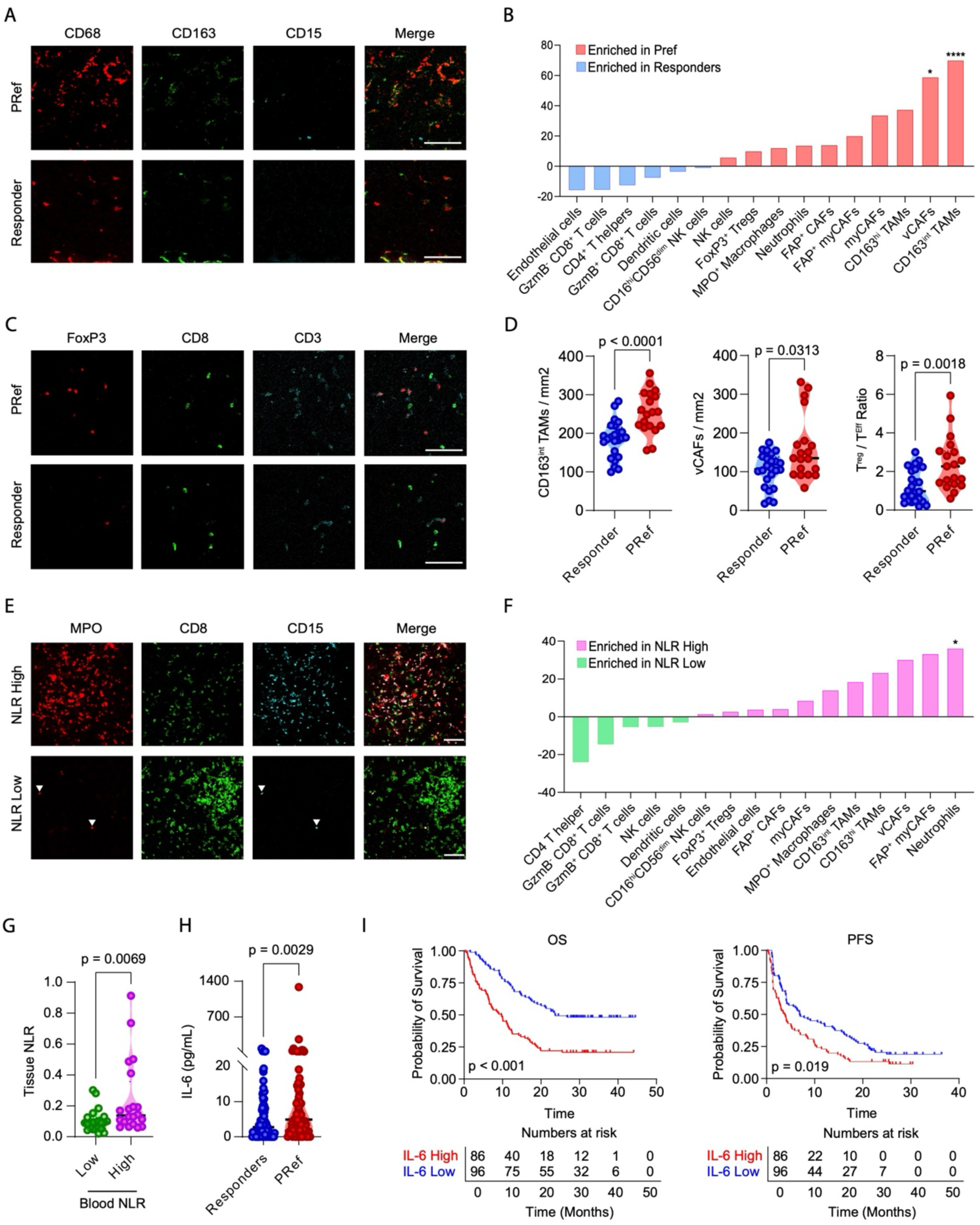
Immune, stromal and systemic predictors to PRef to Atezolizumab plus Bevacizumab. (**A-B**) CD163^Int^ TAMs and vCAFs are strongly enriched in PRef. Figure 3B shows the mean difference of every cell type for PRef vs Responders. (**C-D**) PRef patients possess a higher T^Reg^ / T^Eff^ ratio. (**E-G**) Systemic NLR mimics the intratumour NLR. Figure 3F shows the mean difference of every cell type for NLR High vs NLR Low patients. (**H**) Serum IL-6 expression is higher in PRef than in Responders. (**I**) HCC Patients with high serum IL-6 levels have lower OS and PFS. IL-6, Interleukin 6; PFS, Progression-free survival.

In addition, the ratio between FoxP3^+^ T^reg^ and Granzyme B^+^ CD8^+^ T-cells (GzmB^+^ CD8^+^ T-cells), termed as T^reg^ / T^eff^ ratio, was significantly higher in PRef vs. responders (p<0.01, ***Figure 3C-D***). Cell-cell interaction analysis revealed that PRef patients’ tumours in general are enriched for interactions between anti-tumour immune cells (CD8^+^ T cells, CD4^+^ T helpers, CD16^Hi^ CD56^Dim^ NK cells) and immunosuppressive cells (Neutrophils, CD163^Int^ and CD163^Hi^ TAMs and FoxP3^+^ Tregs). Specifically, PRef tumours were enriched for interactions between Tregs and GzmB^+^CD8^+^ T cells, neutrophils with CD4^+^ T helpers and CD4^+^ T helpers with CD163^Int^ and CD163^Hi^ TAMs (***Supplementary Figure 3A***).

Because primary refractoriness was associated with higher levels of systemic inflammation, we contrasted the characteristics of neutrophil and lymphocyte infiltration within the tumour in patients with NLR ≥3 and <3. Patients with peripheral blood NLR ≥3 harboured tumours with higher neutrophil/lymphocyte infiltration within the HCC TME (p = 0.0069) (***Figure 3E-G***). Cell-cell interaction analysis showed that the tumours of patients with a high NLR were specifically enriched for interactions between anti-tumour immune cells and neutrophils, and not immunosuppressive cells in general like those identified comparing PRef and responders (***Supplementary Figure 3B***).

To further characterise the relationship between PRef and systemic inflammation, we measured a panel of baseline serum cytokines (IL-2, IL-4, IL-6, IL-10, IL-17A, IFNγ, and TNFα) in a subset of Cohort 1 patients (***Supplementary Figure 4***). Among these, IL-6 -a pro-inflammatory cytokine and a well-established marker of resistance to immunotherapy in both tissue and blood^25^ had a higher concentration in PRef versus responders expressed between PRef and responders (p=0.0029) (***Figure 3H*)**. Additionally, patients with higher IL-6 levels had reduced OS and PFS compared to those with lower IL-6 levels (***Figure 3I*)**.

### Immunosuppressive myeloid cell infiltration is a robust hallmark of primary refractoriness to A+B and impairs prognosis

To validate the myeloid cell / T cell imbalance seen in PRef identified in Cohort 1, we used bulk RNAseq data from pre-treatment tumour samples from Cohorts 2 and 3. Demographic and baseline characteristics in the biomarker-evaluable patients were generally consistent with those in the whole population recruited to the GO30140 and IMBrave150 trials (***Supplementary Tables 6)*** and so were clinical outcomes (***Supplementary Figure 5-6***).

We first performed an enrichment cell analysis and evaluated several gene signatures reflective of adaptive and innate immunity in Cohort 2 pre-treatment samples. Cell enrichment analysis on RNASeq datasets confirmed IMC data in portraying PRef as associated with an immunosuppressive, myeloid-enriched landscape characterised by CD4^+^ and CD8^+^ T-cell depletion (***Supplementary Figure 7***).

In an attempt to identify distinctive molecular drivers of PRef, we performed genome-wide differential expression (DEG) analysis in Cohort 2 patients (n=119) as a discovery Cohort and then independently verified in Cohort 3 patients (n=110). We identified differentially expressed genes specifically dysregulated in PRef compared to responders associated with immunosuppression and senescence. These included repression of T-cell effector genes including *IFNG*, *GZMB*, *CD8A* and *FASLG* transcriptional traits predictive of immunotherapy response^26–29^. PRef samples were also characterised by a complex gene expression profile with upregulation of immunosuppressive transcripts involved in myeloid infiltration (i.e. S100A9, CXCL5, NOS2), systemic inflammation and T^reg^ infiltration (i.e. *IL-6, AREG*), evasion from phagocytosis (i.e. *CD24*), stromal activity (i.e. ACTA1), hypoxia and acidosis (i.e. CA12, ATP6V1B1), and microenvironment alteration (i.e. *APOA4*). While responders possessed an enrichment in T and NK cell identity plus activation transcripts (i.e. CD8A, NCR1, NCAM1, NKG7, GZMB, FASLG, TXK, IFNG), antigen presentation (i.e. HLA.DQA1), platelets (ITGA2B), and anti-tumour myeloid activity (CXCL9). These findings were supported by hallmark GSEA, which confirmed PRef samples to harbour a significant enrichment of immunosuppressive pathways including TNF-α, TGF-β, Notch, and signatures related to epithelial-mesenchymal transition, angiogenesis and hypoxia. These features were mirrored by a significant up-regulation in gene expression profiles relating to adaptive immunity such as interferon alpha (IFN-α) and gamma (IFN-γ) in responding patients, to suggest an imbalance in intrinsic tumour immunogenicity across patient groups (***Figure 4A-B***).

**Figure 4.**
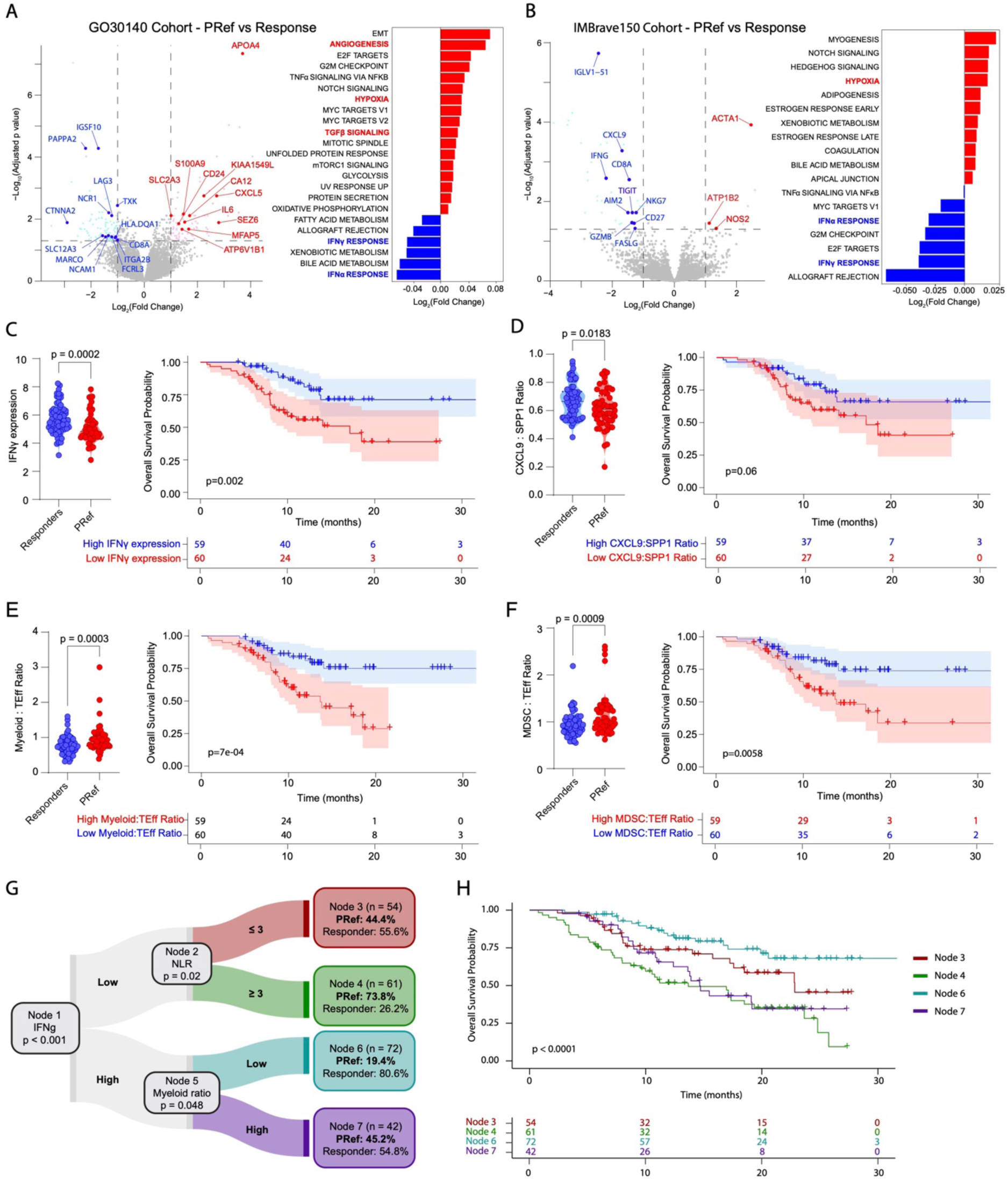
Gene expression profiles of myeloid immunosuppression in PRef. Differential gene expression (volcano plot) and biological pathways (GSEA analysis by CAMERA of the contrast model in the Hallmark gene set collection) comparing PRef and responders in Cohort 2 (**A**) and Cohort 3 (**B**). Expression of INFγ, CXCL9:SPP1 ratio, Myeloid:T^Eff^ ratio and MDSC:T^Eff^ ratio signature plus the respective Kaplan–Meier plots in Cohort 2 stratified by median expression score (**C-F**). Conditional Inference Tree (**G**) and Kaplan-Meier plots for Nodes 3, 4, 6 and 7 (**H**). IFNγ, Interferon γ; CXCL9, C-X-C motif chemokine ligand 9; SPP1, Osteopontin; T^Eff^, T Effector; MDSC, Myeloid Derived Suppressor Cell.

Paired analyses performed in Cohort 3 patients demonstrated that PRef samples were associated with up-regulation of similar immunosuppressive pathways including Notch and hypoxia signalling pathways (***Figure 4B***), validating findings from Cohort 2 RNAseq experiments. Similarly to Cohort 2, several gene sets previously reported to be associated with response to immunotherapy reflective of an inflamed/immune enriched and IFN-γ rich microenvironment were downregulated in PRef and especially expressed in responding patients, supporting the immunosuppressive phenotype typical of PRef (***Figure 4B***).

We then analysed immune signatures and observed a significant reduction in key immune effector populations, including IFN-γ (p<0.001), T^eff^ (p=0.001), NK (p=0.04) and T^cyt^ (p=0.007) signatures (***Figure 4C, Supplementary Figure 8***). While no significant differences were observed in M1/M2 tumour-associated macrophages (TAM) signatures (***Supplementary Figure 8***), we assessed the CXCL9:SPP1 ratio, a recently identified marker of macrophage polarity. We observed a differential expression in our Cohort, with PRef characterised by a low CXCL9:SPP1 ratio (p=0.018) (***Figure 4D***), consistent with hypoxia-driven immune suppression and tumour progression. We also observed a significantly higher myeloid:T^eff^ cells ratio (p=0.0003) and MDSC/Teff ratio (p=0.0009) (***Figure 4E-F***), confirming the IMC data.

### The immune contexture typical of PRef is independently associated with poor patients’ survival

As a further step, we evaluated whether the immune signatures defining PRef were independently associated with patient’s survival. Patients with low expression of the IFN-γ, and T^eff^ signatures, along with high MDSC:T^eff^ and myeloid:T^eff^ cell ratios, had significantly worse OS (IFN-γ signature: HR 2.82, 95%CI 1.42–5.6, p=0.002; T^eff^ signature: HR 2.27, 95%CI 1.16-4.46, p=0.014: MDSC:T^eff^ cells ratio signature: HR 2.54, 95%CI 1.28– 5.04, p=0.006; myeloid:T^eff^ cells ratio signature: HR 3.11, 95% CI 1.56–6.2, p<0.001, ***Figure 4C, E-F; Supplementary Figure 8C; Supplementary Figure 9***).

### Model-based recursive partitioning identifies systemic inflammation and depleted tumour immunogenicity as determinants of primary refractoriness to A+B

To verify the relative contribution of routine pre-treatment variables over the underlying tumour immune contexture as a determinant of PRef to A+B, we integrated clinical and transcriptome data from Cohorts 2 and 3 using a CTree.

We selected NLR and ALBI given their independent prognostic role and combined these with IFN-γ, and T^eff^ signatures, along with the MDSC:T^eff^ and myeloid:T^eff^ cell ratio signatures, based on the prognostic relevance demonstrated before. AFP levels were excluded due to the high proportion (>50%) of missing values in the biomarker-evaluable patients.

This analysis provided a hierarchical disposition of characteristics independently associated with PRef. Notably, repression of IFN-γ related transcripts emerged as the most significant factor associated with PRef, especially when combined with high systemic inflammation, as indicated by an NLR≥3, contributing to shape 73.8% of the risk of PRef in these patients. Conversely, IFN-γ signature overexpression did not entirely preclude PRef, especially in patients with an elevated myeloid:T^eff^ ratio, which doubled the risk compared to those with lower ratios (45% vs. 19.4%, ***Figure 4G***). Consistent with these findings, Kaplan–Meier survival analysis demonstrated significantly different overall survival outcomes across the identified patient subgroups (***Figure 4H***).

## Discussion

In a rapidly expanding treatment landscape, where multiple ICI combinations and TKIs are available for the treatment of unresectable-advanced HCC, early identification and improved mechanistic understanding of primary resistance to treatment represents a clinical imperative to broaden the benefit of anti-cancer immunotherapy in HCC.

Leveraging large, globally accrued patient datasets, including landmark clinical trials of A+B in this indication, our study provides solid and validated criteria to define primary therapeutic resistance to A+B. We found that PRef, defined as the achievement of early disease progression or short-lived HCC stabilisation, affects approximately 45% of patients treated with A+B in clinical trials and routine practice.

This substantial subset of patients exhibits distinctive clinical and immunological features. PRef was more likely in patients with advanced-stage HCC and was associated with worse ALBI grade and elevated NLR. Hypoalbuminemia and high NLR, two recognised biomarkers of systemic inflammation ^30, 31^, were strongly associated with PRef, suggesting an unopposed pro-inflammatory state in these patients. Consistent with these findings, PRef patients exhibited higher intra-tumoural expression and elevated circulating IL-6 levels, a circulating cytokine that controls the switch from acute to chronic inflammation, reinforcing the link between systemic inflammation and an immune microenvironment polarised towards an immune suppressive status ^25^.

Given the strong association between PRef and systemic inflammation, we next examined the TME to dissect the underlying immune landscape associated with treatment failure. Using IMC on pre-treatment biopsies, we found that PRef tumours were characterised by significantly denser myeloid infiltration, T-cell depletion and a more strongly immunosuppressive TME. The presence of a myeloid-dominant phenotype was further and independently validated in clinical trial cohorts, demonstrating that PRef represents a distinct IFN-γ-low, myeloid-enriched immune state.

Among myeloid subsets, TAMs emerged as a key determinant of PRef. Our IMC analysis revealed that PRef tumours exhibited an increased density of CD163^+^ TAMs, including the different spectrums of CD163 expression level, CD163^Hi^ and CD163^Int^ M2-like TAMs, showing the last ones the strongest association with PRef patients. CD163^Int^ TAMs are macrophages that possess both newly recruited and immunomodulatory phenotypes, previously reported to cause chronic liver disease^32^, probably by suppressing T-cell lymphocytes via immune checkpoint axes or secretion of factors like Osteopontin.

To refine our understanding of TAM polarisation, we evaluated CXCL9 and SPP1 (the gene encoding Osteopontin) expression as markers of anti-and pro-tumour macrophage polarisation in our pre-treatment samples. CXCL9 polarisation, induced by IFN-γ, is associated with an anti-tumour immune response, while SPP1 polarisation is upregulated under hypoxic conditions and linked to immune suppression^33^. In our study, whereas not statistically significant, the CXCL9:SPP1 ratio seems to be associated with survival outcomes from A+B, with a low ratio consistently observed in PRef patients. Furthermore, these patients exhibited low IFN-γ signature expression and enrichment of hypoxia-related pathways, reinforcing the link between an immunosuppressive TME and primary resistance.

Beyond individual markers, our findings revealed a broader imbalance in the myeloid compartment of PRef tumours. While NLR has long been recognized as a prognostic indicator in oncology ^22^, our work first described a direct parallel between elevated peripheral NLR and the enrichment of immunosuppressive myeloid cells within the TME. The strong correlation between elevated NLR and intratumoral immune composition suggests a systemic-local immune axis that may contribute to immunotherapy resistance. This finding has the potential for stratifying HCC patients that can benefit from neutrophil-based therapies, currently in clinical trials^34–36^.

Whilst independently associated with prognosis and refractoriness to treatment, the systemic pro-inflammatory response associated with PRef is known to be conditioned by redundant and pleiotropic molecular drivers. For this reason, we opted to integrate findings from our transcriptomic experiments and review of patient data to define the hierarchical relationships among resistance mechanisms. Using CTree we identified IFN-γ repression as the primary determinant of PRef risk, particularly in the presence of elevated systemic inflammation (NLR ≥3). Presence of these two, independent traits defined a high-risk subgroup with a 74% probability of PRef, representing the most unfavourable biological scenario. Notably, even among patients with high IFN-γ expression, an elevated myeloid:T^eff^ cell ratio more than doubled the risk of PRef (45% vs. 19.4%), underscoring the critical role of the balance between effector and suppressive immune populations in shaping treatment outcomes. This hierarchical model not only provides deeper insights into resistance mechanisms but also supports patient stratification based on the dominant resistance pathway, with potential implications for personalized therapeutic strategies.

Several limitations of our study should be acknowledged. Although our multi-cohort approach provides robust validation of determinants and clinical characteristics of patients with PRef to A+B, the retrospective design of our study exposes the risk selection bias, particularly in the observational cohort. In addition, analyses of tissue samples were performed on archival samples, which may not fully reflect the dynamic changes in the TME at the time of treatment initiation. Whilst convergent, IMC and transcriptomic analyses were conducted on different patient subsets with non-overlapping experimental readouts, potentially limiting direct cross-comparisons between these datasets. While we identified key biological features associated with PRef, the causal relationships among these factors remain to be fully elucidated. Lastly, while our conditional inference tree model offers a framework for patient stratification, prospective validation in independent cohorts is necessary before clinical implementation.

Despite these limitations, our findings have significant implications for drug development and study design in HCC. The prominence of myeloid-mediated immunosuppression suggests that targeting this axis could enhance the efficacy of A+B. Several emerging therapeutic strategies aim to counteract myeloid-driven resistance by modulating macrophage and myeloid cell function within the tumour microenvironment. Among these, combining A+B with anti-T-cell immunoreceptor with Ig and ITIM domains (TIGIT) inhibitors (i.e. tiragolumab) has shown promise, leveraging both T-cell activation and myeloid modulation to enhance anti-tumour immunity^37, 38^. In patients with NSCLC, high intratumoral T^reg^ and myeloid infiltration correlated with improved benefit from atezolizumab plus tiragolumab versus atezolizumab alone^39^, suggesting preferential activity in tumours that share immunobiologic characteristics with PRef HCC. Other strategies to overcome myeloid-driven tumour immune escape may involve blocking CD24, another hallmark of PRef that suppresses macrophage phagocytosis via Siglec-10^40, 41^, LILRB2 (ILT4), known to transmit inhibitory signals in macrophages^42^ (NCT04524871), or use of CSF1R inhibitors, which deplete or reprogramme immunosuppressive TAMs ^43^, as well as therapies targeting the SIRPα-CD47 axis, which enhance phagocytosis ^44^. Our hierarchical model suggests that patients with repressed IFN-γ signalling and NLR ≥3, who have a 74% probability of PRef, may represent prime candidates for myeloid-targeting combination strategies.

In conclusion, our study provides the first comprehensive characterisation of PRef to A+B in HCC, establishing PRef as a distinct biological entity defined by systemic inflammation, myeloid cell infiltration, and T-cell depletion. These findings not only enhance our understanding of immunotherapy resistance in HCC but also lay the groundwork for biomarker-driven combination strategies aimed at overcoming primary resistance. Future studies should focus on the prospective validation of these findings, elucidation of resistance evolution during treatment, and evaluation of targeted interventions to enhance immunotherapy efficacy in resistant patient populations.

## Data availability

All clinical, raw RNA-seq data for the GO30140 and IMbrave150 trials are deposited in the European Genome-Phenome Archive under accession no. <u>EGAS00001005503</u>. Qualified researchers may request access to individual patient-level data through the clinical study data request platform (https://vivli.org/). Further details on Roche’s criteria for eligible studies are available at https://vivli.org/members/ourmembers. For further details on Roche’s Global Policy on the Sharing of Clinical Information and how to request access to related clinical study documents, see https://www.roche.com/research_and_development/who_we_are_how_we_work/clinical_trials/our_commitment_to_data_sharing.htm.

## Funding

This research was funded by the National Research Foundation of Korea (NRF) grants, supported by the Korean government (MSIT): grant numbers NRF 2023R1A2C2004339 to H.J.C.

J.L. is supported by an Academy of Medical Sciences Springboard award (SBF009\1103), Royal Society Research Grant (RG\R2\232323), JGW Patterson Foundation primer grant, CRUK program grants (DRCRPG-Nov22/100007) and an MRC project grant (MR/Y003365/1). E.R.G funded by W.E. Harker PhD studentship (C0260N3038). J.L. and E.R.G are supported by a NIHR BRC project grant (NIHR203309).

## Supporting information

Supplementary Methods and Results

## Acknowledge.

The authors would like to acknowledge the following sources of funding: European Society of Medical Oncology (ESMO) Translational Fellowship (to PL).

## Notes

### Competing Interest Statement

The authors have declared no competing interest.

