## Supplementary Figures and Tables for "Clinical and molecular characterisation of primary refractoriness to atezolizumab plus bevacizumab in patients with unresectable hepatocellular carcinoma"

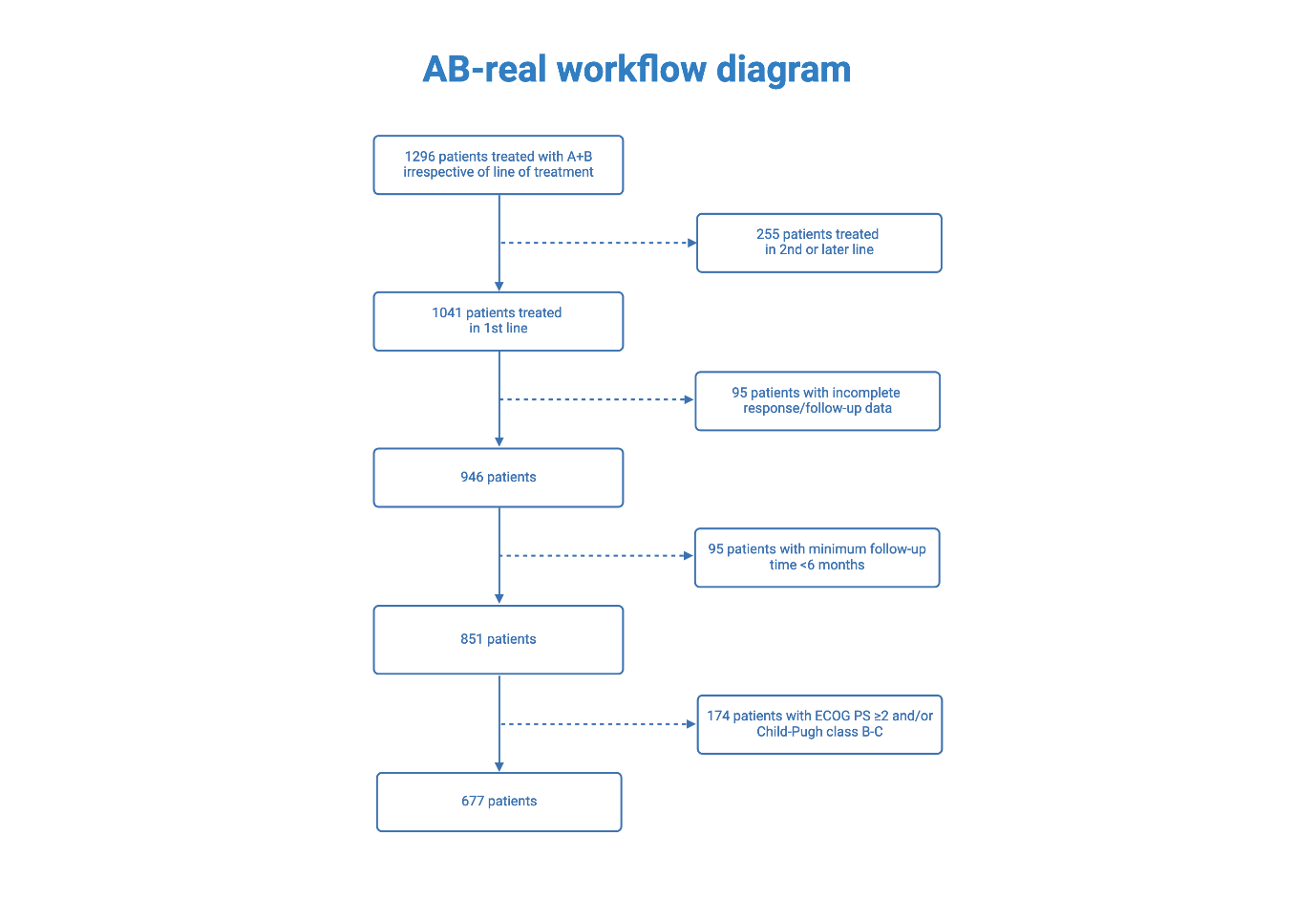

**Supplementary Figure 1a -  Flowchart of the patients included in the clinical analysis for Cohort 1**

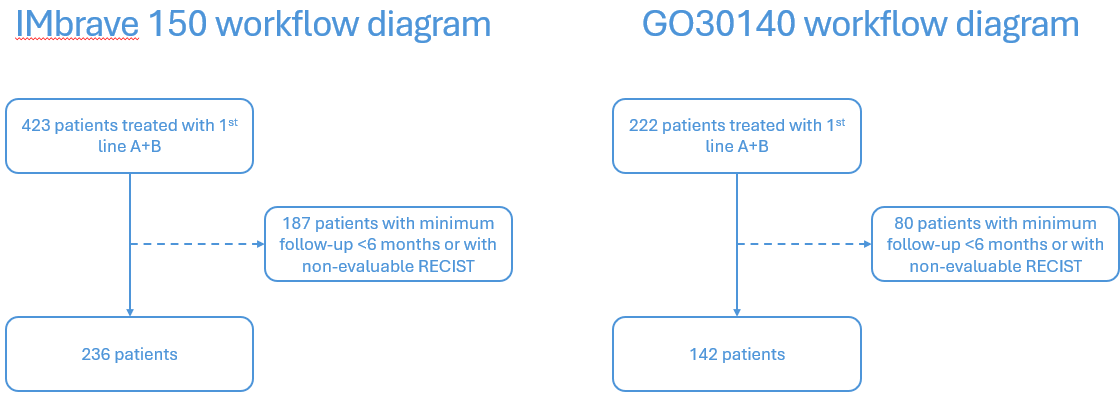

**Supplementary Figure 1b -  Flowchart of the patients included in the clinical analysis for Cohort 2 and 3**

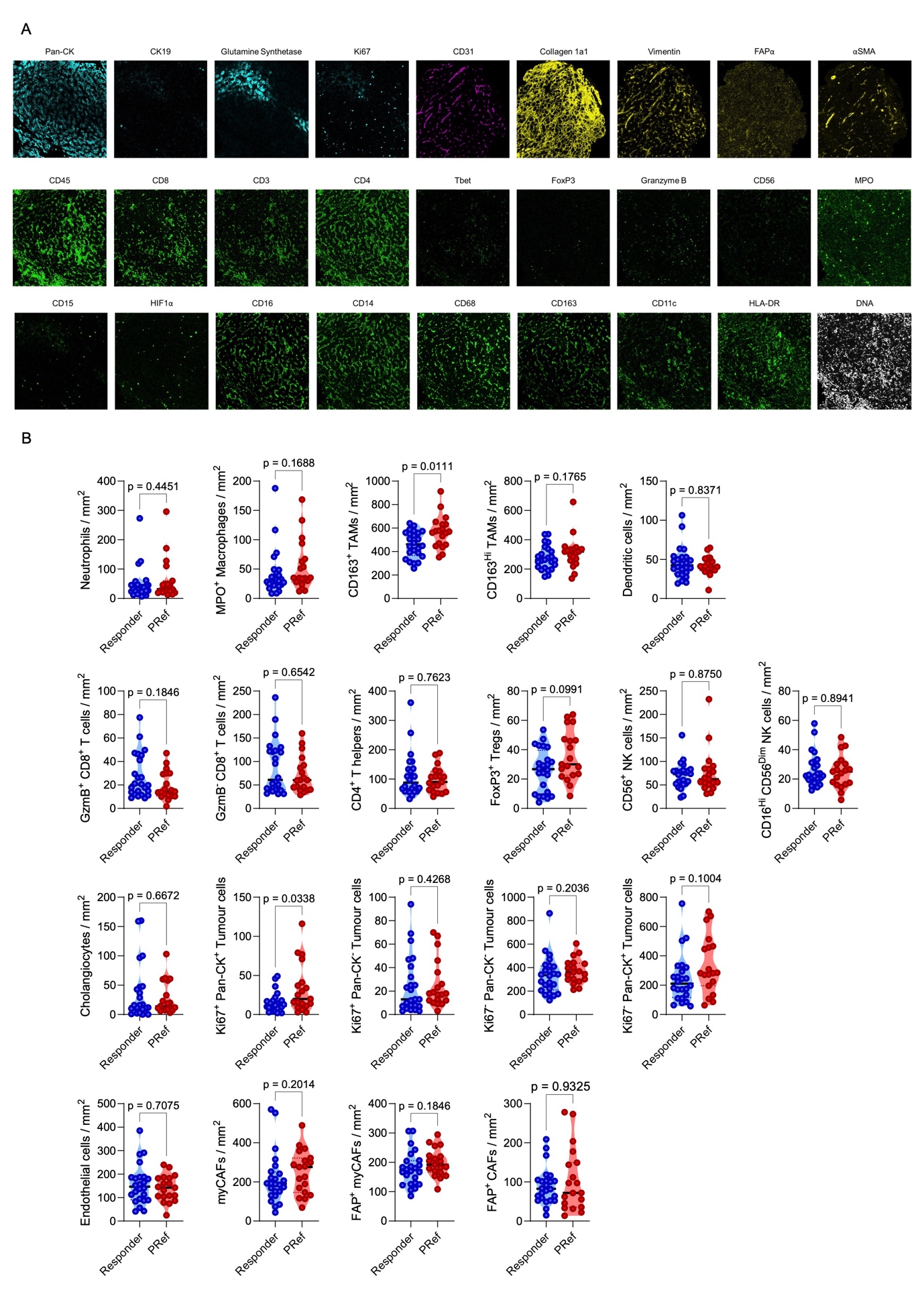

**Supplementary figure 2. IMC Marker Panel and Cell densities for PRef and Responders.** (**A**) Single stains for each marker of the IMC panel, (**B**) Cell density for every cell type of the HCC microenvironment between PRef and Responders.

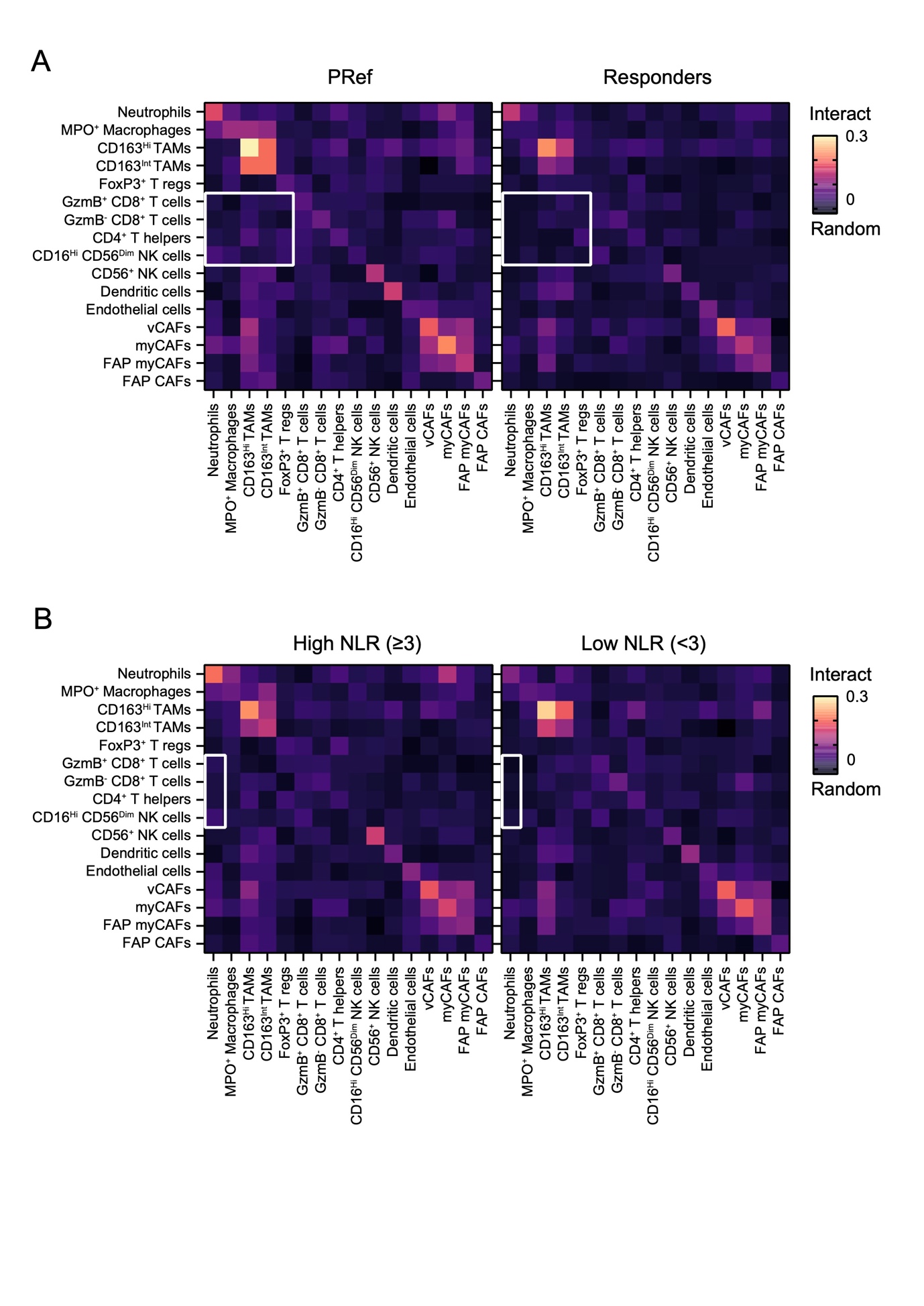

**Supplementary figure 3. Cell-cell interactions in Pref vs Responder and NLR High vs NLR Low.** (**A**) Cellular interactions in PRef and Responder HCC patients (**B**) Cellular interactions in patients with High vs Low NLR.

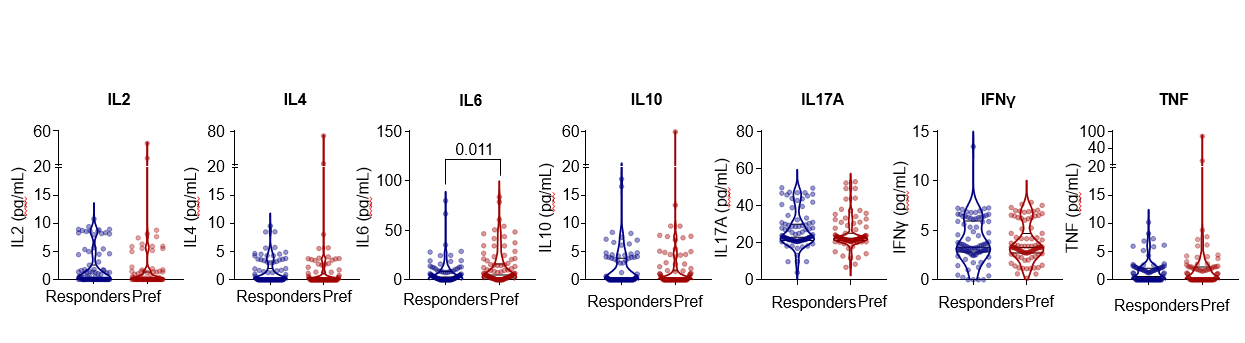

**Supplementary Figure 4** **– Cytokine analysis in Cohort 1**. Baseline serum cytokines (IL-2, IL-4, IL-6, IL-10, IL-17A, IFNγ, and TNFα) in a subset of Cohort 1 patients

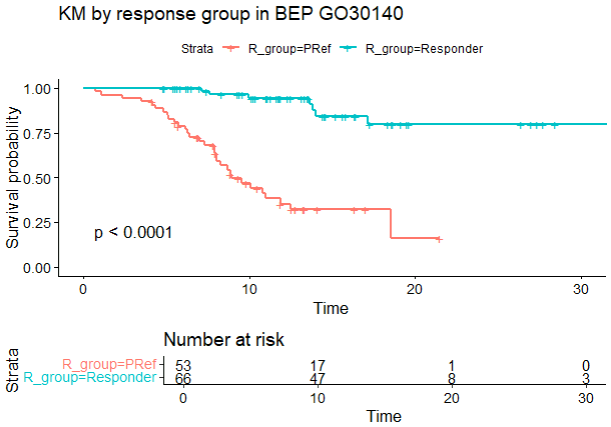

**Supplementary Figure 5** – OS in PRef and Responder in biomarker evaluable population (BEP) in Cohort 2

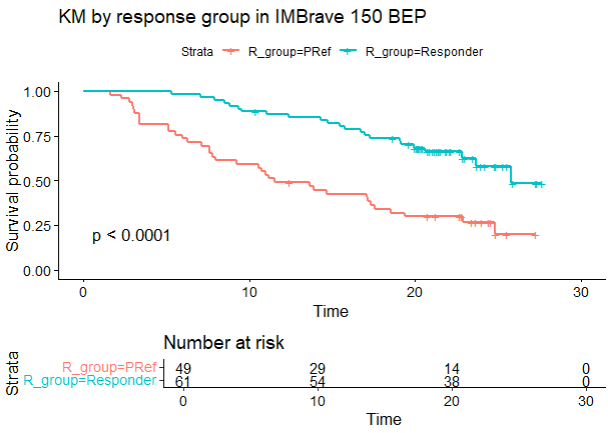

**Supplementary Figure 6 – OS in PRef and Responder in biomarker evaluable population** (BEP) in Cohort 3

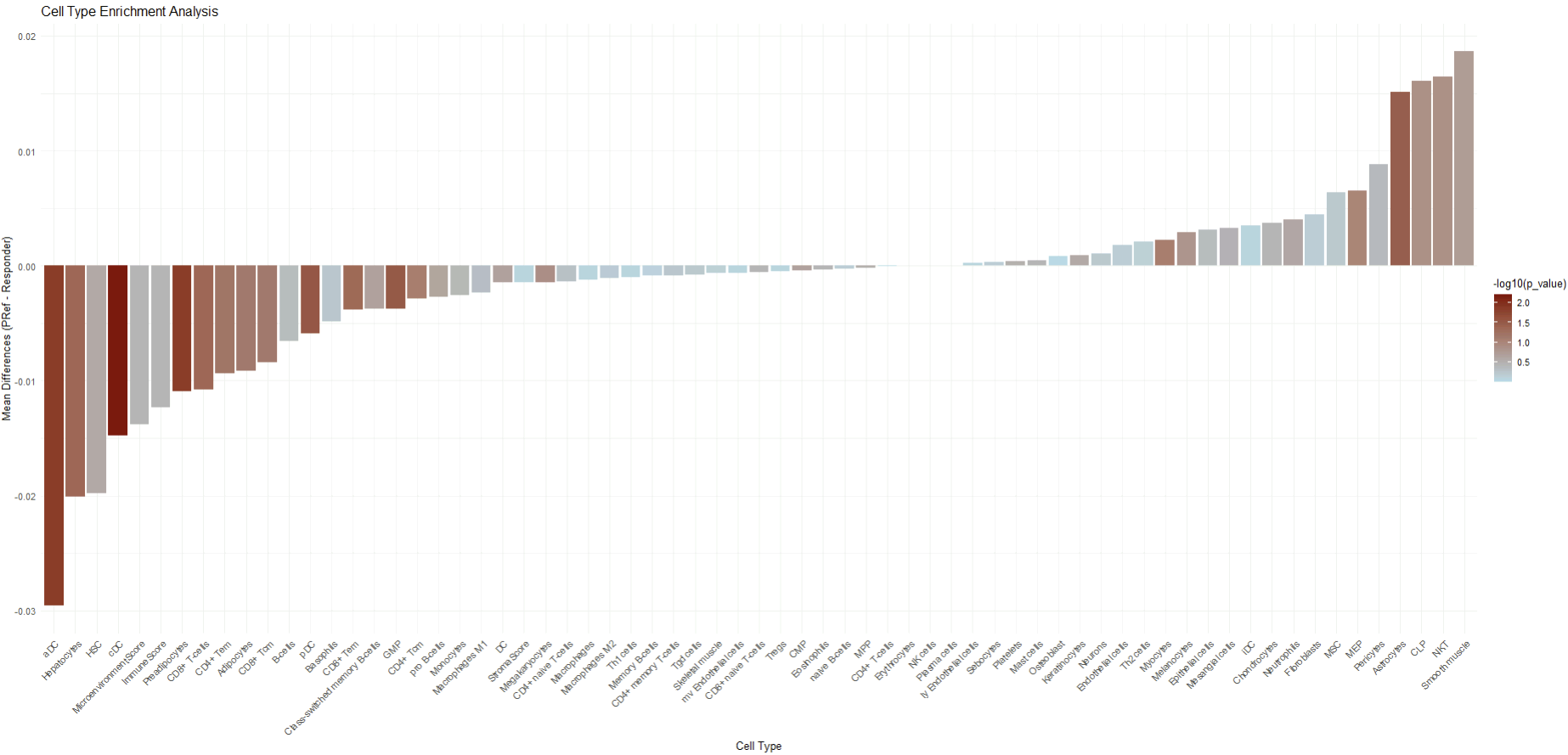

**Supplementary Figure 7 – XCell deconvolution in Cohort 2**

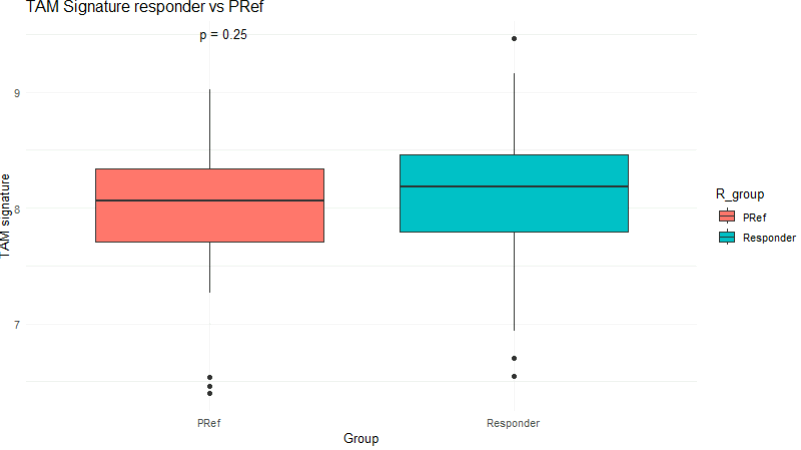

**Supplementary Figure 8a - Expression of TAM signature**

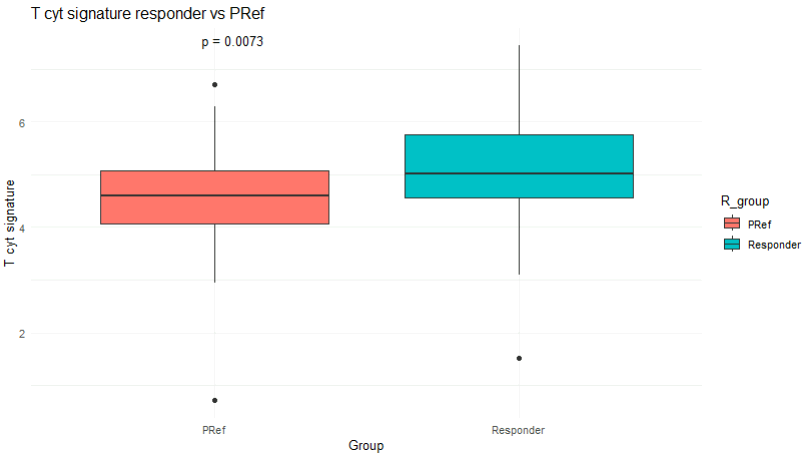

**Supplementary Figure 8b - Expression of T cytotoxic signature**

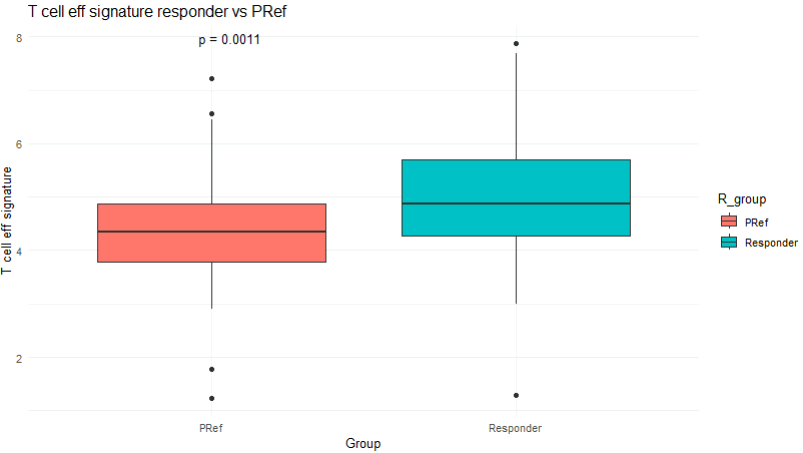

**Supplementary Figure 8c - Expression of T effector signature**

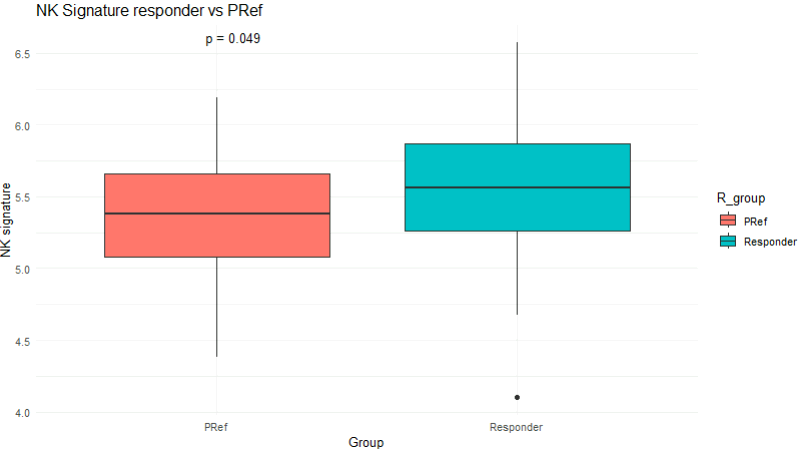

**Supplementary Figure 8d - Expression of NK signature**

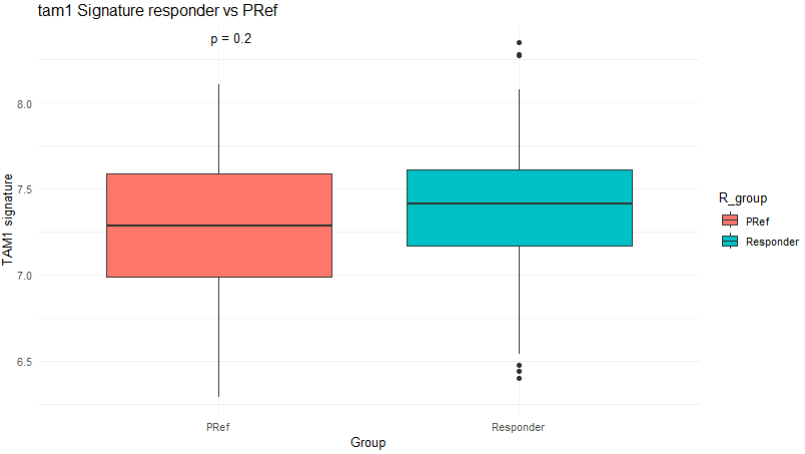

**Supplementary Figure 8e - Expression of TAM M1 signature**

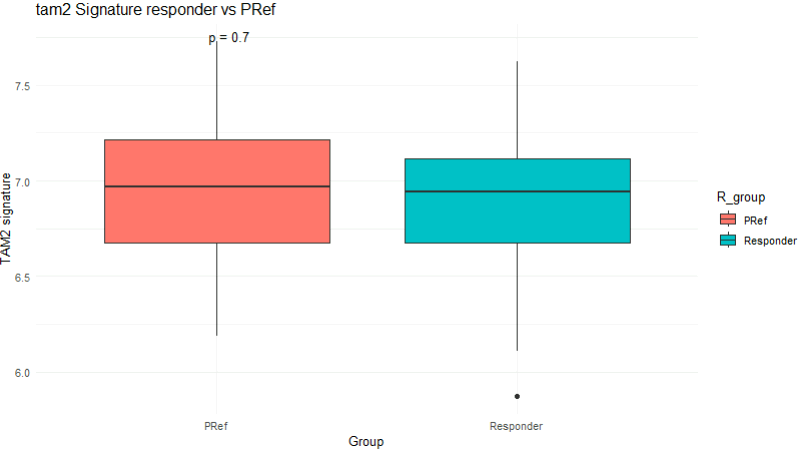

**Supplementary Figure 8f - Expression of TAM M2 signature**

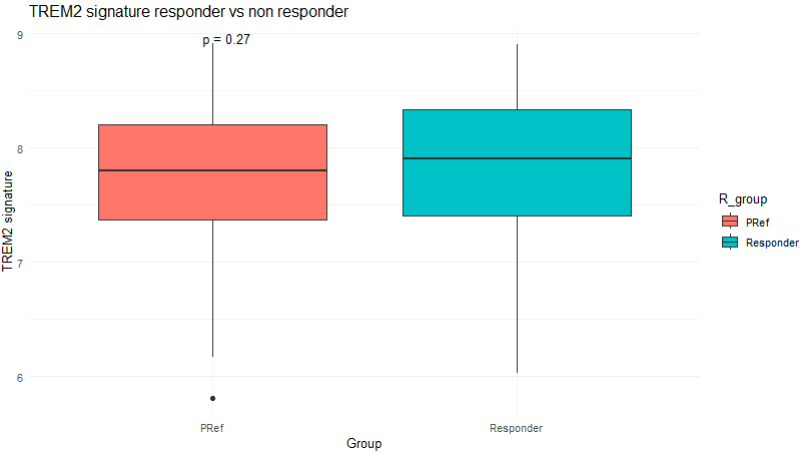

**Supplementary Figure 8g - Expression of TREM2 signature**

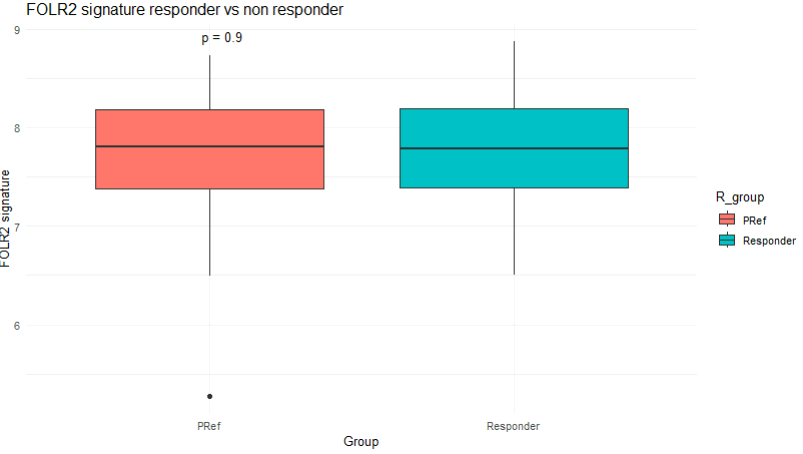

**Supplementary Figure 8h - Expression of TAM FOLR2+ signature**

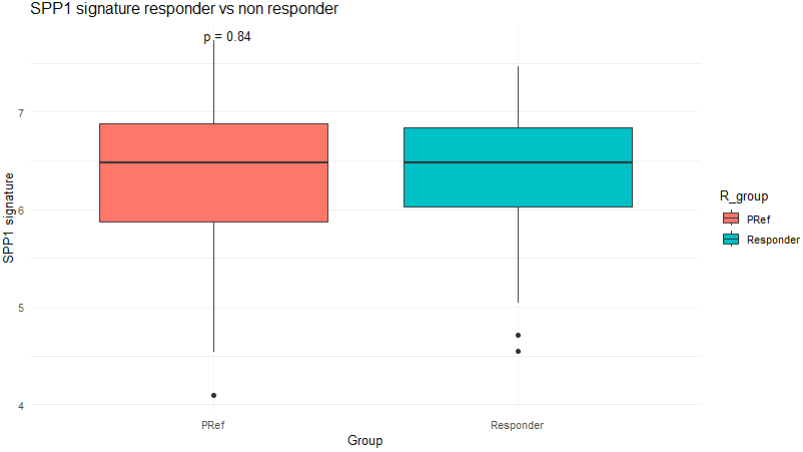

**Supplementary Figure 8i - Expression of TAM SPP1+ signature**

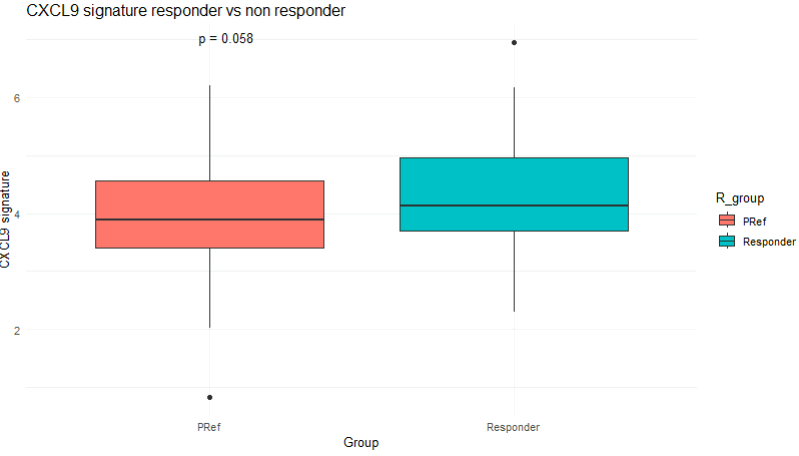

**Supplementary Figure 8l - Expression of TAM CXCL9+ signature**

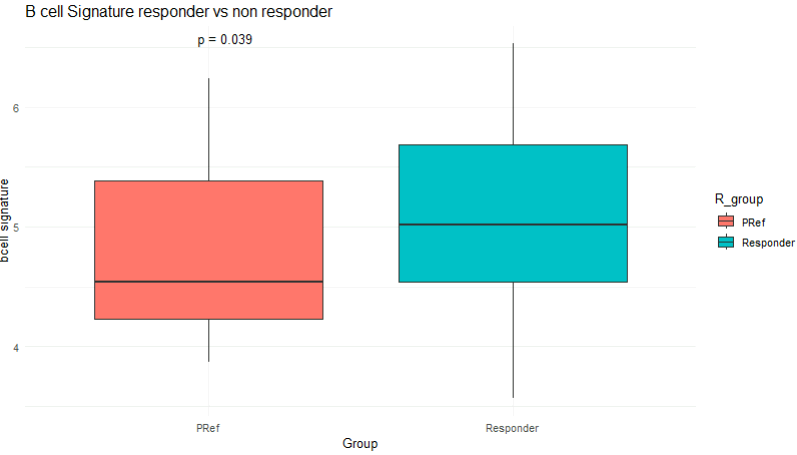

**Supplementary Figure 8m - Expression of B cell signature**

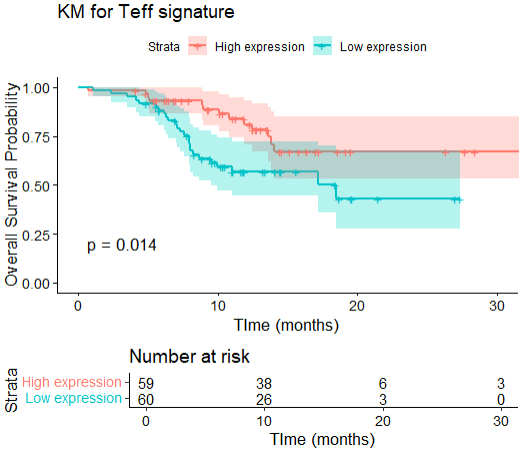

**Supplementary Figure 9 - Kaplan–Meier plots in Cohort 2 stratified by median expression of Teff signature score**

**Supplementary Table 1. IMC Antibodies.** IMC Antibody clones, Catalogue number and Suppliers. * Antibodies marked with an asterisk passed the QC for IHC specificity, but were excluded from the analysis because of a suboptimal staining by IMC.

| **Metal**  **isotope** | **Marker** | **Working**  **dilution** | **Clone** | **Catalogue**  **number** | **Supplier** |
| --- | --- | --- | --- | --- | --- |
| 141 Pr | MPO | 1:100 | Polyclonal | AF3667 | R&D |
| 142 Nd | Vimentin | 1:500 | D21H3 | 46173SF | CST |
| 143 Nd | CK19 | 1:100 | EP1580Y | ab195872 | Abcam |
| 144 Nd | CD15 | 1:50 | W6D3 | 323002 | BioLegend |
| 145 Nd | CD163 | 1:100 | EDHu-1 | MCA1853 | BioRad |
| 146 Nd | Ki67 | 1:100 | D2H10 | 44092SF | CST |
| 147 Sm | Pan-CK | 1:100 | AE1+AE3 | CF190321 | ThermoFisher |
| 148 Nd | Glut. Synthetase | 1:500 | GT1055 | MA5-27749 | ThermoFisher |
| 149 Sm | HLA-DR | 1:100 | LN3 | 14-9956-82 | ThermoFisher |
| 150 Nd | Col1a1 | 1:500 | E8F4L | 81375SF | CST |
| 151 Eu | CD8α | 1:100 | C8/144B | 372902 | BioLegend |
| 152 Sm | CXCR2***** | 1:50 | EPR22301-103 | ab245982 | Abcam |
| 153 Eu | CD4 | 1:100 | EPR6855 | ab181724 | Abcam |
| 154 Sm | XCR1 | 1:100 | D2F8T | 63684SF | CST |
| 156 Gd | δTCR | 1:250 | H-41 | sc-100289 | Santa Cruz B |
| 157 Gd | MYH11***** | 1:500 | SP314 | ab240983 | Abcam |
| 158 Gd | CD31 | 1:500 | 89C2 | 85873SF | CST |
| 159 Tb | CD45RO***** | 1:200 | UCHL1 | 304202 | BioLegend |
| 160 Gd | Tbet | 1:1000 | D6N8B | 27112SF | CST |
| 161 Dy | CD3ε | 1:500 | D7A6E™ | 85061BF | CST |
| 162 Dy | CD79a | 1:100 | EP3618 | ab239891 | Abcam |
| 163 Dy | CD14 | 1:100 | Polyclonal | 17000-1-AP | ProteinTech |
| 164 Dy | PD-L1***** | 1:500 | SP142 | ab236238 | Abcam |
| 165 Ho | CD45 | 1:200 | D9M8I | 47937SF | CST |
| 166 Er | FAPα | 1:100 | Polyclonal | AF3716 | R&D |
| 167 Er | FoxP3 | 1:750 | 236A/E7 | ab96048 | Abcam |
| 168 Er | CD56 | 1:100 | E7X9M | 88856SF | CST |
| 169 Tm | CD11c | 1:100 | D3V1E | 93233SF | CST |
| 170 Er | CD16 | 1:100 | EPR16784 | ab215977 | Abcam |
| 171 Yb | Granzyme B | 1:100 | D6E9W | 79903SF | CST |
| 173 Yb | CK18 | 1:500 | Polyclonal | PA5-14263 | ThermoFisher |
| 174 Yb | αSMA | 1:200 | 1A4 | 14-9760-82 | ThermoFisher |
| 175 Lu | HIF1α | 1:500 | EP1215Y | ab210073 | Abcam |
| 176 Yb | CD68 | 1:100 | C68/684 | ab213098 | Abcam |

**Supplementary Table 2 - Curated signatures representing potential clinically relevant pathways and immune subsets**

| Signature | Gene Contente | Source |
| --- | --- | --- |
| B cell | AKNA, ARHGAP25, CCL21, CD1D, CD2, CD27, CD37, CD38, CLEC17A, CLEC9A, CLECL1, FAIM3, FAM65B, GIMAP4, MAP4K1, PAX5, TNFRSF17, TRAF3IP3, BANK1, CD22, CYBB, ETS1, FAIM2C, FCGR1A, FCGR2B, FCRL3, FCRL5, FCRLA, HLA-DQ9, HLA-DOB, HLA-DOA1, HVCN1, KIAA0226, NCF1, PTPRC, P2RY10, PINC, SPI00, STAT1, TAGAP, TXNIP, ZCCHC7, ACAD4, CCNA2, CDKN3, CLCN5, ENPP1, FCER1A, FCRL4, MRC, NEDD1, RCL1, SOX5, STAT5A, STAT5B, TLR7 | Angelova M., Genome Biology, 2015 |
| Interferon gamma response | IDO1, CXCL10, CXCL9, HLA-DRA, STAT1, IFNG | Ayers M., *JCI Insight*, 2017 |
| Myeloid inflammation | CXCL1, CXCL2, CXCL3, CXCL8, IL6, PTGS1 | McDermott D.F., *Nature Medicine*, 2018 |
| Myeloid-derived-suppressor cell (MDSC) | FCN1, S100A6, VCAN, S100A8, S100A9, S100A12 | Zhang Q., *Cell*, 2019 |
| Natural killer (NK) cell | AKT3, AXL, CDH2, CRTAM, CYTH1, FASLG, GRB2, KLRG1, LILRB5, LST1, MAP4K4, NOTCH3, PIK3CG, PILRA, PLCG2, SIGLEC7, SIGLEC9 | Angelova M., *Genome Biology*, 2015 |
| T effector cell (Teff) | CXCL9, GZMB, PRF1 | McDermott D.F., *Nature Medicine*, 2018 |
| T regulatory cell (Treg) | FOXP3, CD4, CTSC, TNFRSF4, FOXP3, TNFRSF18, IKZF2, IL2RA | Zhang Q., *Cell*, 2019 |
| T cytotoxic cell (T Cyt) | GZMA, PRF1 | Rooney MS, Cell, 2015 |
| Tumor-associated macrophage (TAM) | C1QA, C1QB, APOE, TREM2, GPNMB, SLC40A1 | Zhang Q., *Cell*, 2019 |
| TAM M1 | C1QA, C1QB, SPP1, LGMN, APOC1, CTSD, CD68, MS4A4A, PDL3, FOLR2, GPNMB, CD63, CTSB, PSAP, FLT1, CD14, LIPA, TYROBP, NPC2, DAB2, FCGRT, RNASE1, CTSC, FCER1G, SLC40A1, APOE, CCL3, CCL3L3, MS4A7, CCL4L2, CD5L, VCAM1, CST3, SDC3, ITM2B, CXCL4, CXCL2, HES1, FOS, IER3 | Sharma M, Cell, 2020 |
| TAM M2 | RNASE1, CTSD, CTSB, NUPR1, FLT1, APOC1, GPNMB, APOE, CTSZ, LGALS1, PLTP, FABP5, TREM2, CXCL3, COLEC12, HSPA1A, PLIN2, SDC1, CTSL, HSPA1B, MMP19, PLD3, ABL2, CEBPB | Sharma M, Cell, 2020 |
| Macrophages SPP1+ | SPP1, C1QB, LPL, MS4A4A, VSIG4, SDSL | Bill R, Science, 2023 |
| Macrophages CXCL9+ | CXCL9, CCL18, CHIT1, CMKLR1, GAL3ST4, RAB42 | Bill R, Science, 2023 |
| Macrophages TREM2+ | TREM2, C1QA, C1QB, C1QC, APOE, SPP1 | Li H, Front Immunol., 2023 |
| Macrophages FOLR2+ | SEPP1, SPP1, LGMN, RNASE1, FOLR2 | Xiang C, Cell Death Dis., 2023 |

**Supplementary Table 3 - Baseline Characteristics for Cohort 1 according to SITC Criteria**

|  | **Whole population**  N = 677^1^ | **Pref**  N = 313^1^ (46%) | **Responders**  N = 364^1^ (54%) | **p-value** |
| --- | --- | --- | --- | --- |
| **Sex, n (%)** |  |  |  |  |
| Male | 530 (78%) | 241 (77%) | 289 (79%) | 0.5 |
| Female | 147 (22%) | 72 (23%) | 75 (21%) |  |
| **Age at treatment start, median (IQR)** | 67 (59 – 75) | 67 (59 - 75) | 68 (59 - 75) | 0.8 |
| **Etiology, n (%)** |  |  |  |  |
| Viral | 460 (62%) | 194 (62%) | 226 (62%) | >0.9 |
| Non-viral | 257 (38%) | 119 (38%) | 138 (38%) |  |
| **Portal Vein Thrombosis, n (%)** |  |  |  |  |
| Yes | 202 (30%) | 113 (36%) | 89 (24%) | **0.001** |
| No | 475 (70%) | 200 (64%) | 275 (76%) |  |
| **Extra-hepatic Spread, n (%)** |  |  |  |  |
| Yes | 313 (46%) | 155 (49.6%) | 158 (43.5%) | 0.13 |
| No | 364 (54%) | 158 (50.4%) | 206 (56.5%) |  |
| **ECOG PS, n (%)** |  |  |  |  |
| 0 | 350 (52%) | 149 (48%) | 201 (56%) |  |
| 1 | 322 (46%) | 161 (52%) | 161 (44%) | 0.064 |
| Missing | 5 | 3 | 2 |  |
| **Child Pugh score, n (%)** |  |  |  |  |
| A5 | 418 (64%) | 170 (55%) | 248 (73%) |  |
| A6 | 231 (36%) | 137 (45%) | 94 (27%) | **<0.001** |
| Missing | 28 | 6 | 22 |  |
| **BCLC stage, n (%)** |  |  |  |  |
| A+B | 164 (24%) | 62 (20%) | 102 (28%) |  |
| C | 509 (76%) | 249 (80%) | 260 (72%) | **0.0017** |
| Missing | 4 | 2 | 2 |  |
| **ALBI grade, n (%)** |  |  |  |  |
| 1 | 320 (47.5%) | 119 (38%) | 201 (56%) |  |
| 2 + 3 | 353 (52.5%) | 192 (62%) | 161 (44%) | **<0.001** |
| Missing | 4 | 2 | 2 |  |
| **NLR, n (%)** |  |  |  |  |
| < 3 | 410 (60%) | 167 (37%) | 243 (60%) |  |
| ≥ 3 | 214 (32%) | 124 (63%) | 90 (40%) | **<0.001** |
| Missing | 54 (8%) | 22 | 31 |  |
| **AFP, n (%)** |  |  |  |  |
| <400 | 257 (38%) | 165 (54%) | 244 (68%) |  |
| ≥400 | 409 (60%) | 141 (46%) | 116 (32%) | **<0.001** |
| Missing | 11 (2%) | 7 | 4 |  |
| ^1^Median (Q1 - Q3); n (%) |  |  |  |  |

**Supplementary Table 4 - Baseline Characteristics for Cohort 2-3 according to SITC Criteria**

|  | **Whole population**  N = 378 | **PRef**  N = 168^1^ (44%) | **Responders**  N = 210^1^ (56%) | **p-value** |
| --- | --- | --- | --- | --- |
| **Sex, n (%)** |  |  |  |  |
| Male | 310 (82%) | 139 (83%) | 171 (81%) | 0.846 |
| Female | 68 (18%) | 29 (17%) | 39 (55%) |  |
| **Age at treatment start, median (IQR)** | 63 (56 – 71) | 61 (55 - 69) | 64 (59 - 72) | **0.007** |
| **Etiology, n (%)** |  |  |  |  |
| Viral | 246 (65%) | 118 (70%) | 146 (70%) | 0.155 |
| Non-viral | 114 (35%) | 50 (30%) | 64 (30%) |  |
| **Portal Vein Thrombosis, n (%)** |  |  |  |  |
| Yes | 35 (10%) | 21 (13%) | 14 (7%) | 0.078 |
| No | 342 (90%) | 146 (87%) | 196 (93%) |  |
| **Extra-hepatic Spread, n (%)** |  |  |  |  |
| Yes | 247 (65%) | 123 (73%) | 124 (59%) | **0.006** |
| No | 132 (35%) | 46 (27%) | 86 (41%) |  |
| **ECOG PS, n (%)** |  |  |  |  |
| 0 | 227 (60%) | 84 (50%) | 143 (68%) |  |
| 1 | 151 (40%) | 84 (50%) | 67 (32%) | **0.001** |
| **Child Pugh score, n (%)** |  |  |  |  |
| A5 | 269 (71%) | 113 (67%) | 156 (74.5%) |  |
| A6 | 102 (27%) | 50 (30%) | 52 (25%) | 0.251 |
| B7 | 5 (2%) | 4 (2%) | 1 (0.5) |  |
| **BCLC stage, n (%)** |  |  |  |  |
| A+B | 47 (12%) | 13 (7.7%) | 44 (21%) |  |
| C | 321 (88%) | 155 (92.3%) | 166 (79%) | **0.001** |
| **ALBI grade, n (%)** |  |  |  |  |
| 1 | 158 (42%) | 64 (38%) | 94 (45%) |  |
| 2 + 3 | 220 (58%) | 104 (62%) | 116 (55%) | 0.230 |
| **NLR, n (%)** |  |  |  |  |
| < 3 | 208 (55%) | 70 (42%) | 138 (66%) |  |
| ≥ 3 | 170 (45%) | 98 (58%) | 72 (34%) | **<0.001** |
| **AFP, n (%)** |  |  |  |  |
| <400 | 181 (48%) | 74 (44%) | 108 (51%) |  |
| ≥400 | 108 (28%) | 54(32%) | 54 (24%) | 0.178 |
| Missing | 92 (24%) | 40 (24%) | 52 (25%) |  |
| ^1^Median (Q1 - Q3); n (%) |  |  |  |  |

**Supplementary Table 5 - Multiple Cox Regression for OS with manual selection of covariates**

|  | **Cohort 1** | | **Cohorts 2-3** | |
| --- | --- | --- | --- | --- |
| **Covariates** | **Hazard Ratio (95% CI)** | **p-value** | **Hazard Ratio (95% CI)** | **p-value** |
| **Time-dependent response according to SITC Criteria (Yes vs No)** | 4 (2.8-5.5) | **<0.001** | 4.9 (3.6–6.7) | **<0.001** |
| **NLR (≥3 vs <3)** | 1.37 (1.10-1.72) | **0.004** | 1.54 (1.12-2.1) | **0.007** |
| **AFP (≥ 400 vs <400)** | 1.67 (1.34-2.07) | **<0.001** | 1.83 (1.3-2.61) | **<0.001** |
| **Viral aethiology (Yes vs No)** | 1.00 (0.79-1.25) | 0.97 | 1.32 (0.95-1.83) | 0.09 |
| **ALBI Grade (1 vs 2-3)** | 1.57 (1.27-1.96) | **<0.001** | 2.09 (1.5-2.93) | **<0.001** |
| **PVT (Yes vs No)** | 1.06 (0.84-1.34) | 0.59 | 1.31 (0.79-2.18) | 0.29 |
| **ECOG PS (1-2 vs 0)** | 1.20 (0.97-1.50) | 0.09 | 1.02(0.75-1.38) | 0.91 |
| **EHS (Yes vs No)** | 1.01 (0.81-1.25) | 0.94 | 1.08 (0.77-1.5) | 0.65 |

**Cohort 1**

Concordance= 0.749 (se = 0.015)

Likelihood ratio test= 153 on 8 df, p<0.001

Wald test= 128.2 on 8 df, p<0.001

Score (logrank) test= 140.6 on 8 df, p<0.001

**Cohorts 2-3**

Concordance= 0.729 (se = 0.02)

Likelihood ratio test= 115.9 on 11 df, p<0.001

Wald test= 105.9 on 11 df, p<0.001

Score (logrank) test= 116.5 on 11 df, p<0.001

**Supplementary Table 6 - Baseline Characteristics for BEP Cohort 2-3**

|  | Overall |
| --- | --- |
| n | 229 |
| Age (median [IQR]) | 63.00 [57.00, 71.00] |
| NLR (median [IQR]) | 2.69 [1.94, 3.96] |
| Sex (%) |  |
| Female | 46 (20.1) |
| Male | 183 (79.9) |
| Race (%) |  |
| Asian | 146 (63.8) |
| Black or african american | 10 (4.4) |
| Unknown | 5 (2.2) |
| White | 68 (29.7) |
| ECOG, n(%) |  |
| 0 | 139 (60.7) |
| 1 | 90 (39.3) |
| Child Pugh score, n (%) |  |
| A5 | 169 (73.8) |
| A6 | 55 (24.0) |
| B7 | 3 (1.3) |
| AFP, n (%) |  |
| <400 ng/mL | 95 (41.5) |
| ≥400 ng/mL | 53 (23.1) |
| NA | 81 (35.4) |
| Etiology, n (%) |  |
| Hepatitis B | 115 (50.2) |
| Hepatitis C | 46 (20.1) |
| Non-viral | 68 (29.7) |
| Stage, n (%) |  |
| IB | 2 (0.9) |
| II | 24 (10.5) |
| IIIA | 22 (9.6) |
| IIIB | 32 (14.0) |
| IVA | 21 (9.2) |
| IVB | 128 (55.9) |
| BCLC stage, n (%) |  |
| A-B | 32 (14.0) |
| C | 197 (86.0) |
| Extra-hepatic Spread, n (%) |  |
| No | 80 (34.9) |
| Yes | 149 (65.1) |
| Portal Vein Thrombosis, n (%) |  |
| NA | 1 (0.4) |
| No | 216 (94.3) |
| Yes | 12 (5.2) |
| NLR, n (%) |  |
| ≥3 | 104 (45.4) |
| <3 | 125 (54.6) |
| ALBI, n (%) |  |
| 1 | 101 (44.1) |
| 2-3 | 128 (55.9) |
