## Supplementary Methods and Results for "Clinical and molecular characterisation of primary refractoriness to atezolizumab plus bevacizumab in patients with unresectable hepatocellular carcinoma"

**Study design and patients.**

The AB-real Cohort is a prospectively maintained database including HCC patients treated with A+B across 23 tertiary referral centres in Europe, North America, and Asia accrued between May 15, 2018, and September 1, 2024.

The study was conducted according to the ethics guidelines in the Declaration of Helsinki. For the HCC Cohort, ethical approval to conduct this study was granted following review of the study protocol by the Imperial College Tissue Bank (Reference Number R16008) and locally by the ethical committee of each participating site.

GO30140 (NCT02715531) is an open-label, multicenter, multi-arm, phase Ib whose primary analysis has been previously reported^1^. The study included five Cohorts, with the two HCC Cohorts (groups A and F). In group A, all patients included had received A+B. In group F, patients were randomly assigned (1:1) to receive A+B or atezolizumab alone. Primary endpoints were (1) confirmed overall response rate (ORR) in all patients who received the combination treatment for group A and (2) progression-free survival (PFS) in the intention to treat (ITT) population in group F.

IMbrave150 (NCT03434379) is a global, open-label, phase III trial testing A+B against sorafenib in unresectable HCC^2^ . The co-primary endpoints were OS and PFS in the ITT population, as assessed by an IRF according to RECIST 1.1.

For GO30140 and IMbrave150 studies, the protocols were approved by the ethics committee or institutional review board at each centre. Inclusion and exclusion criteria are described in detail in previously published papers^1, 2^.

**Main inclusion criteria**

Main inclusion criteria included: age at least 18 years old; Eastern Cooperative Oncology Group (ECOG) Performance Status (PS) ≤ 2; histological or radiological diagnosis of HCC in accordance with the American Association for the Study of Liver Diseases (AASLD) criteria^3^; diagnosis of aHCC disease as per Barcelona Clinic Liver Cancer (BCLC) criteria^4^- BCLC C or BCLC B not amenable to locoregional therapy; atezolizumab plus bevacizumab as first-line systemic treatment; patient must be evaluable by RECIST v1.1. Patients with Child-Pugh class B or C cirrhosis and patients who received combination treatments including anti-CTLA-4 were excluded.

### **Treatment and safety**

Cohort 1.

The treatment regimen was approved in every center after a multidisciplinary evaluation and according to local clinical practice. All the patients received a first line treatment with the combination of atezolizumab 1200 mg and bevacizumab 15 mg/Kg every three weeks. Treatment was maintained until loss of clinical benefit or in the presence of unacceptable toxicity. Data regarding patients’ demographics and clinical status were collected retrospectively and prospectively maintained and updated at each participating site. Treatment related adverse events (trAEs) were managed according to drug data sheet. In accordance with the updated analysis of IMbrave150, we included in the safety and efficacy analysis all patients who received at least one dose of atezolizumab-bevacizumab in the first line setting. TrAEs were assessed at every contact with the patient and were graded according to the National Cancer Institute Common Terminology Criteria for Adverse Events (CTCAE) v5.0. Principal investigators at each site had at least 5 years of expertise in administering systemic anticancer treatment.

Cohort 2

 In group A and in the combination therapy group in group F, patients were given 1200 mg atezolizumab intravenously and 15 mg/kg bevacizumab intravenously every 3 weeks. Patients in the atezolizumab monotherapy group in group F received 1200 mg atezolizumab intravenously every 3 weeks. Treatment continued until loss of clinical benefit or unacceptable toxicity. Dose reductions were not permitted.

Cohort 3

In this open-label, phase 3 trial, patients were randomly assigned in a 2:1 ratio to receive atezolizumab plus bevacizumab or sorafenib.

Patients assigned to the atezolizumab–bevacizumab group received 1200 mg of atezolizumab plus 15 mg per kilogram of body weight of bevacizumab intravenously every 3 weeks; patients assigned to the sorafenib group received 400 mg of sorafenib orally twice daily. Patients received their assigned drugs until unacceptable toxic effects occurred or there was loss of clinical benefit.

**Assessments and radiological-response outcomes**

Cohort 1

Radiological response to treatment was assessed according to RECIST criteria v1.1 every 9-12 weeks as per clinical practice. Treatment duration was calculated from the date of the first dose of atezolizumab plus bevacizumab to the date the treatment discontinuation. OS was defined as the time from the date of the first dose of the treatment to the date of death. PFS was defined as the time from the date of the first dose of the treatment to the date of radiological evidence of tumor progression or the date of death, whichever occurred first. Radiological response and radiological diagnosis of progression were assessed locally by experienced radiologists in each center, without any central imaging review.

Cohort 2

Tumour assessments were done every 8 weeks for the first year and every 12 weeks thereafter by the investigator until patient death, disease progression, or initiation of further systemic cancer therapy. A confirmed objective response was one that was observed on two consecutive occasions at least 4 weeks apart. Patients could receive treatment beyond progression, according to RECIST 1.1, if there was evidence of investigator-assessed clinical benefit.

Cohort 3

Tumors were assessed by computed tomography or magnetic resonance imaging at baseline and every 6 weeks until week 54 and then every 9 weeks thereafter.

**Machine-learning approach for tumour infiltrating lymphocytes (TILs) identification and quantification**

We applied a supervised machine learning (ML) algorithm to 37 and 184 H&E of Cohort 1 and Cohort 2-3, respectively, developed using QuPath (v.0.5.0) and followed the previously described procedure with some modifications^5, 6^. In summary, the analysis began with color deconvolution to estimate stain vectors and normalize the RGB channels on a per-slide basis. Cell segmentation was then performed using the “WatershedCellDetection” plugin, optimized for hematoxylin-based nuclear detection. Detection was carried out on the "Hematoxylin OD" channel with a pixel size of 0.5 µm. Background smoothing included a background radius of 8.0 µm, a median filter with a 1.0 µm radius and Gaussian smoothing with a sigma of 1.5 µm. Cells were identified using a threshold of 0.1 and refined using a watershed post-processing step with a maximum background value of 2. Detected objects were filtered by area, retaining only those between 7 µm² and 500 µm². Each nucleus was expanded by 2 µm to approximate the full cell boundary. Nuclei were included in the final segmentation, boundaries were smoothed and cellular measurements were recorded for downstream classification. The quality of cell detection was evaluated by a pathologist before proceeding to classification.

Object/cell classification was conducted using QuPath built-in random forest classifier, configured with the following parameters: max_depth=25, min_samples=5, max_trees=100 and active_vars=10. During model development, additional features were incorporated including RGB vectors, hematoxylin and eosin optical density channels, multi-scale sigma features (1, 2, 4 and 8) and difference-of-Gaussian–derived metrics (Laplacian, weighted deviation and gradient magnitude). Training data consisted of regions of interest (512×512 pixels) extracted from 71 images comprising a total of 5324 annotated cells (3319 tumor cells, 1119 tumor-infiltrating lymphocytes (TILs) and 886 stroma/other cells). TILs per mm^2^ were defined as mononuclear immune cells including lymphocytes and plasma cells^7, 8^.

The classifier was finalized following qualitative review by a pathologist and then applied to tumor tissue cores, excluding areas with artifacts or necrosis, across whole slide images. Tumor cores were defined as regions containing tumor epithelium and intra-tumoral stroma, excluding extra-tumoral stroma and adjacent normal tissue. In cases where multiple tumor cores were present, all available cores were used for classification per patient.

**Tissue gene expression analysis.**

Bulk RNA sequencing (RNAseq) was performed using TruSeq RNA Access technology (Illumina) as previously described^9^. Following total RNA purification, RNAseq reads were first aligned to ribosomal RNA sequences to remove ribosomal reads. The remaining reads were aligned to the human reference genome (NCBI Build 38) using GSNAP v.2013-11-01. To quantify gene expression levels, the number of reads mapped to the exons of each RefSeq gene was calculated using the functionality provided by the R/Bioconductor package GenomicAlignments.

**Statistical analysis.**

Statistical analyses were performed using R v.4.3.1. We estimated survival curves using the Kaplan-Meier method and compared them with the log-rank test. Hazard ratio (HR) estimates and 95% CI and P values were determined using the stratified Cox proportional hazards model and log-likelihood test. Proportionality assumption was validated by confirming the absence of a significant correlation between event time and Schoenfeld residual. Multivariate adjustment to HR was performed by the addition of sex, age, Eastern Cooperative Oncology Group (ECOG) performance status, disease aetiology (viral/nonviral), presence of either macrovascular invasion or extrahepatic spread and level of serum AFP >400 ng/ml, ALBI score and neutrophile-to-lymphocyte ratio (NLR) as terms to the Cox proportional hazards model. ALBI score was obtained using the formula (log10 bilirubin in μmol/l x 0.66) + (albumin in g/l x -0.085). According to the obtained score, the ALBI grade was assigned: for scores ≤-2.60 grade 1, for scores > -2.6 and ≤-1.39 grade 2, for scores > -1.39 grade 3. NLR was obtained dividing the absolute number of neutrophils count (expressed in cells/μl) by the absolute number of lymphocytes count.

To assess the association between SITC-defined response and survival outcomes while minimizing immortal time bias, we employed a time-dependent Cox proportional hazards model. Response status was treated as a time-varying variable, meaning patients with SD were initially classified as PRef. Only those who remained progression-free beyond six months were reclassified as responders, whereas patients with progression or SD for less than six months remained PRef. By dynamically updating response status, this approach enhanced the accuracy of hazard estimates and effectively prevented artificial survival advantages in the responder group. The proportional hazards assumption was assessed using Schoenfeld residuals.

We performed differential gene expression using the R package DESeq2, contrasting between PRef and responders. We defined significant differential expression as absolute log2 fold-change packages>1 and adjusted P < 0.05.

We performed xCell deconvolution analysis on baseline RNA-seq count data for GO30140 group A using the xCell R package. Cell type enrichment scores of each cell type were then compared between PRef and responders using Student’s t-test and assessed for significance using Benjamini–Hochberg-adjusted, two-sided P values.

We calculated gene signature scores for each sample as the arithmetic mean of log2(CPM) expression of all genes in a given signature. GSEA was performed using the MultiGSEA package v.1.1.99 with default parameters, and the Hallmark gene set collection from the molecular signatures database collection (MSigDB) using a differential expression linear model as the input and the CAMERA method for statistics to adjust P values to account for multiple hypothesis testing^10^.

To provide a hierarchical understanding of the determinants of PRef, we implemented a conditional inference tree (CTree) analysis using the PRef state as the endpoint. Variables included in the CTree analysis were selected based on their impact on OS and data completeness. Data processing was performed using the party R package, with an alpha level of 0.05 (corresponding to mincriterion = 0.95). Patients with missing values in any of the included covariates were excluded from the analysis. The results were visualized using Sankey diagrams to illustrate the hierarchical structure of determinants influencing PRef.

Biomarker-evaluable population was analysed separately in both Cohorts 2 and 3 to mitigate batch effects in the RNA-seq data. However, they were combined during the construction of a conditional inference tree model, where the RNA signature was reduced to categorical variables.

**Survival Outcomes**

After a median follow-up of 20.9 (Cohort 1) and 14.3 months (Cohort 2-3), the median OS was 14.9 (9.5-24.9) and 14.4 (8.9 – 25.6) months in Cohort 1 and Cohort 2-3 respectively.
